# Site-specific effects of neurosteroids on GABA_A_ receptor activation and desensitization

**DOI:** 10.1101/2020.04.27.063404

**Authors:** Yusuke Sugasawa, Wayland W. L. Cheng, John R. Bracamontes, Zi-Wei Chen, Lei Wang, Allison L. Germann, Spencer R. Pierce, Thomas C. Senneff, Kathiresan Krishnan, David E. Reichert, Douglas F. Covey, Gustav Akk, Alex S. Evers

**Affiliations:** Departments of Anesthesiology, Washington University in St. Louis, St. Louis, Missouri 63110; Departments of Developmental Biology, Washington University in St. Louis, St. Louis, Missouri 63110; Departments of Radiology, Washington University in St. Louis, St. Louis, Missouri 63110; Departments of Psychiatry, Washington University in St. Louis, St. Louis, Missouri 63110; Departments of Taylor Family Institute for Innovative Psychiatric Research, Washington University in St. Louis, St. Louis, Missouri 63110

**Author notes:** To whom correspondence should be addressed: Professor Alex S. Evers, Department of Anesthesiology, Washington University School of Medicine, Campus Box 8054, St. Louis, Missouri 63110.

**Keywords:** neurosteroids, GABA_A_ receptor, desensitization, radioligand binding, photoaffinity labeling, site-directed mutagenesis

## Abstract

This study examines how site-specific binding to the three identified neurosteroid binding sites in the α_1_β_3_ GABA_A_ receptor (GABA_A_R) contributes to neurosteroid allosteric modulation. We found that the potentiating neurosteroid, allopregnanolone, but not its inhibitory 3β-epimer epi-allopregnanolone, binds to the canonical β_3_(+)–α_1_(-) intersubunit site that mediates receptor activation by neurosteroids. In contrast, both allopregnanolone and epi-allopregnanolone bind to intrasubunit sites in the β_3_ subunit, promoting receptor desensitization and the α_1_ subunit promoting ligand-specific effects. Two neurosteroid analogues with diazirine moieties replacing the 3-hydroxyl (KK148 and KK150) bind to all three sites, but do not potentiate GABA_A_R currents. KK148 is a desensitizing agent, whereas KK150 is devoid of allosteric activity. These compounds provide potential chemical scaffolds for site-specific and general neurosteroid antagonists. Collectively, these data show that differential occupancy and efficacy at three discrete neurosteroid binding sites determine whether a neurosteroid has potentiating, inhibitory, or competitive antagonist activity on GABA_A_Rs.

## INTRODUCTION

Neurosteroids (NS) are endogenous modulators of brain development and function and are important mediators of mood (1-4). Exogenously administered NS analogues have been clinically used as anesthetics and anti-depressants and have therapeutic potential as anti-epileptics, neuroprotective agents and cognitive enhancers (1,2,5-9). The principal target of NS is the γ-aminobutyric acid type A receptor (GABA_A_R). NS can either activate or inhibit GABA_A_Rs. Positive allosteric modulatory NS (PAM-NS) such as allopregnanolone (3α5αP) potentiate the effect of GABA on GABA_A_R currents at low concentrations and directly activate the receptors at higher concentrations (5,10-12). Negative allosteric modulatory NS (NAM-NS), such as epi-allopregnanolone (3β5αP) or pregnenolone sulfate (PS) inhibit GABA_A_R currents (13-17). In addition to enhancing channel opening, PAM-NS increase the affinity of the GABA_A_R for orthosteric ligand binding, an effect thought to be mechanistically linked to channel gating (11,18).

GABA_A_Rs are pentameric ligand-gated ion channels (pLGIC) composed of two α-subunits (α_1–6_), two β-subunits (β_1–3_) and one additional subunit (γ_1–3_, δ, ε, θ or π) (19-21). Each subunit is composed of a large extracellular domain (ECD), a transmembrane domain (TMD) formed by four membrane-spanning helices (TM1-4), a long intracellular loop between TM3 and TM4, and a short extracellular C-terminus (5,19,20,22). NS modulate GABA_A_Rs by binding to sites within the TMDs (1,2,5,6,11,23-28). Specifically, the α subunit TMDs are essential to the actions of PAM-NS (11,23,24,27,28). Mutagenesis studies in α_1_β_2_γ_2_ GABA_A_Rs have identified several residues in the α_1_ subunit, notably Q242 and W246 in TM1, as critical to NS potentiation of GABA-elicited currents (25,26,29). Crystallographic studies have subsequently shown that, in homo-pentameric chimeric receptors in which the TMDs are derived from either α_1_ (24,27) or α_5_ subunits (23), the NS 3α,21dihydroxy-5α-pregnan-20-one (3α5α-THDOC), pregnanolone and alphaxalone bind in a cleft between the α subunits, with the C3-hydroxyl substituent of the steroids interacting directly with Q242 in the α subunit (αQ242). PAM-NS activate these chimeric receptors, and their action is blocked by αQ242L and αQ242W mutations. These studies posit a single canonical intersubunit binding site for NS action that is conserved across the six α subunit isoforms (23,24,27).

An alternative body of evidence suggests that PAM-NS modulation of GABA_A_R function is mediated by multiple mechanisms and/or binding sites. Site-directed mutagenesis has identified multiple disparate residues on GABA_A_Rs that affect NS-induced activation, suggestive of two neurosteroid binding sites; one site mediating potentiation of GABA responses and the other mediating direct activation (25,26). Single channel electrophysiological studies (5,10,30) as well as studies examining neurosteroid modulation of [^35^S]t-butylbicyclophosphorothionate (TBPS) binding (31), have also identified multiple distinct effects of NS, with various structural analogues producing some or all of these effects, consistent with multiple NS binding sites (25,26). Our recent photolabeling studies have confirmed that there are multiple PAM-NS binding sites on α_1_β_3_ GABA_A_Rs (11). In addition to the canonical site at the interface between the TMDs of adjacent subunits (intersubunit site) (11,23,24,27), we identified NS binding sites within the α-helical bundles of both the α_1_ and β_3_ subunits (intrasubunit sites) of α_1_β_3_ GABA_A_Rs (11). 3α5αP binds to all three sites (11); mutagenesis of these sites suggests that the intersubunit and α_1_ intrasubunit sites, but not the β_3_ intrasubunit site, contribute to 3α5αP PAM activity (11). A functional effect for NS binding to the β_3_ intrasubunit site has not been identified.

The 3α-hydroxyl (3α-OH) group is critical to NS activation of GABA_A_Rs and 3β-OH NS lack PAM activity (5,14). Indeed, many 3β-OH NS are GABA_A_R NAMs (14,16). While molecular docking studies have suggested that the 3β-OH NS epi-pregnanolone competes for binding with PAM-NS (23), 3β-OH NS are non-competitive inhibitors with respect to GABA and 3α-OH NS, indicating that they are unlikely to act at the canonical PAM binding site (5,14). Steroids with a sulfate rather than a hydroxyl at the 3-carbon are also GABA_A_R NAMs thought to act at sites distinct from GABA_A_R PAMs (5,13,14,17,32). The precise location of this site is unclear, but crystallographic studies have demonstrated a possible binding site between TM3 and TM4 on the intracellular end of the α-subunit TMD (17,24). While 3β-OH NS and PS both inhibit GABA_A_Rs, they likely act via interactions with distinct sites (5,13,14,16,17,23,24).

The goal of the current study was to determine the specific sites underlying the PAM and NAM actions of NS. We hypothesized that various NS analogues preferentially bind to one or more of the three NS binding sites in the α_1_β_3_ GABA_A_R, stabilizing distinct conformational states (i.e. resting, open or desensitized). To achieve this goal, we used two endogenous NS, the PAM-NS 3α5αP and the NAM-NS 3β5αP and two NS analogues, KK148 and KK150, in which a diazirine replaced the function-critical 3-OH group (33). We examined site-specific NS binding and effects using NS photolabeling (28,34-36) and measurements of channel gating and orthosteric ligand binding. The NS lacking a 3α-OH were devoid of PAM-NS activity, but surprisingly, KK148 and 3β5αP enhanced the affinity of [^3^H]muscimol binding. We interpret this finding as evidence that these compounds preferentially bind to and stabilize desensitized receptors, since both open and desensitized GABA_A_R exhibit enhanced orthosteric ligand binding affinity (37).

The results show that 3α5αP binds to the canonical β(+)–α(-) intersubunit site, stabilizing the open state of the receptor, whereas the 3-diazirinyl NS (KK148 and KK150) bind to this site but do not promote channel opening, and 3β5αP does not occupy this site. These data indicate that NS binding to the intersubunit sites is largely responsible for PAM activity and that the 3α-OH is critical for NS activation. In contrast, 3α5αP, 3β5αP and the 3-diazirinyl NS all bind to both the α_1_ and β_3_ intrasubunit sites. Occupancy of the intrasubunit sites by 3α5αP, 3β5αP and KK148 promotes receptor desensitization. KK150 occupies all three NS binding sites on α_1_β_3_ GABA_A_Rs, but produces minimal functional effect suggesting a possible scaffold for a general NS antagonist. These results shed new light on the mechanisms of NS allosteric modulation of channel function, and demonstrate a novel pharmacology in which related ligands bind to different subsets of functional sites on the same protein, each in a state-dependent manner, with the actions at these sites summating to produce a net physiological effect.

## RESULTS

### Distinct patterns of NS potentiation and enhancement of muscimol binding

The endogenous NS, 3α5αP is known to potentiate GABA-elicited currents (Figure 1A) and enhance [^3^H]muscimol binding to α_1_β_3_ GABA_A_Rs (Figure 1E) (11,18). We examined a series of NS analogues with different stereochemistries or substituents in the 3- and 17-positions: 3β5αP, KK148, and KK150 (structures shown in Figures 1B-D) for their ability to potentiate GABA-elicited currents and enhance orthosteric agonist ([^3^H]muscimol) binding. 3β5αP is the 3β-epimer of 3α5αP. KK148 and KK150 are NS analogue photolabeling reagents, which have a 3-diazirinyl moiety instead of the 3-OH, and differ from each other by the stereochemistry of the 17-ether linkage (33). We observed a discrepancy between the ability of these compounds to potentiate GABA-elicited currents and their ability to enhance [^3^H]muscimol binding in α_1_β_3_ GABA_A_Rs. None of the NS analogues lacking a 3α-OH potentiated GABA-elicited currents (Figures 1B-D). However, both 3β5αP and KK148 significantly enhanced [^3^H]muscimol binding (Figure 1E). KK150, in contrast, did not potentiate GABA-elicited currents and minimally enhanced [^3^H]muscimol binding (Figures 1D-E). Collectively, these data show that, NS analogues with different stereochemistry or substituents at the 3- and 17-positions show distinct patterns in modulation of α_1_β_3_ GABA_A_R currents and orthosteric ligand binding. We hypothesized that these patterns are a consequence of the various NS analogues stabilizing distinct conformational states of the GABA_A_R, possibly by binding and acting at different sites. Notably, the compounds with a 3-OH (3α5αP, 3β5αP) are ten-fold more potent than those with a 3-diazirine (KK148, KK150) in enhancing [^3^H]muscimol binding (Figure 1E), suggesting that the 3-OH is an important determinant of binding affinity to the site(s) mediating these effects.

**FIGURE 1:**
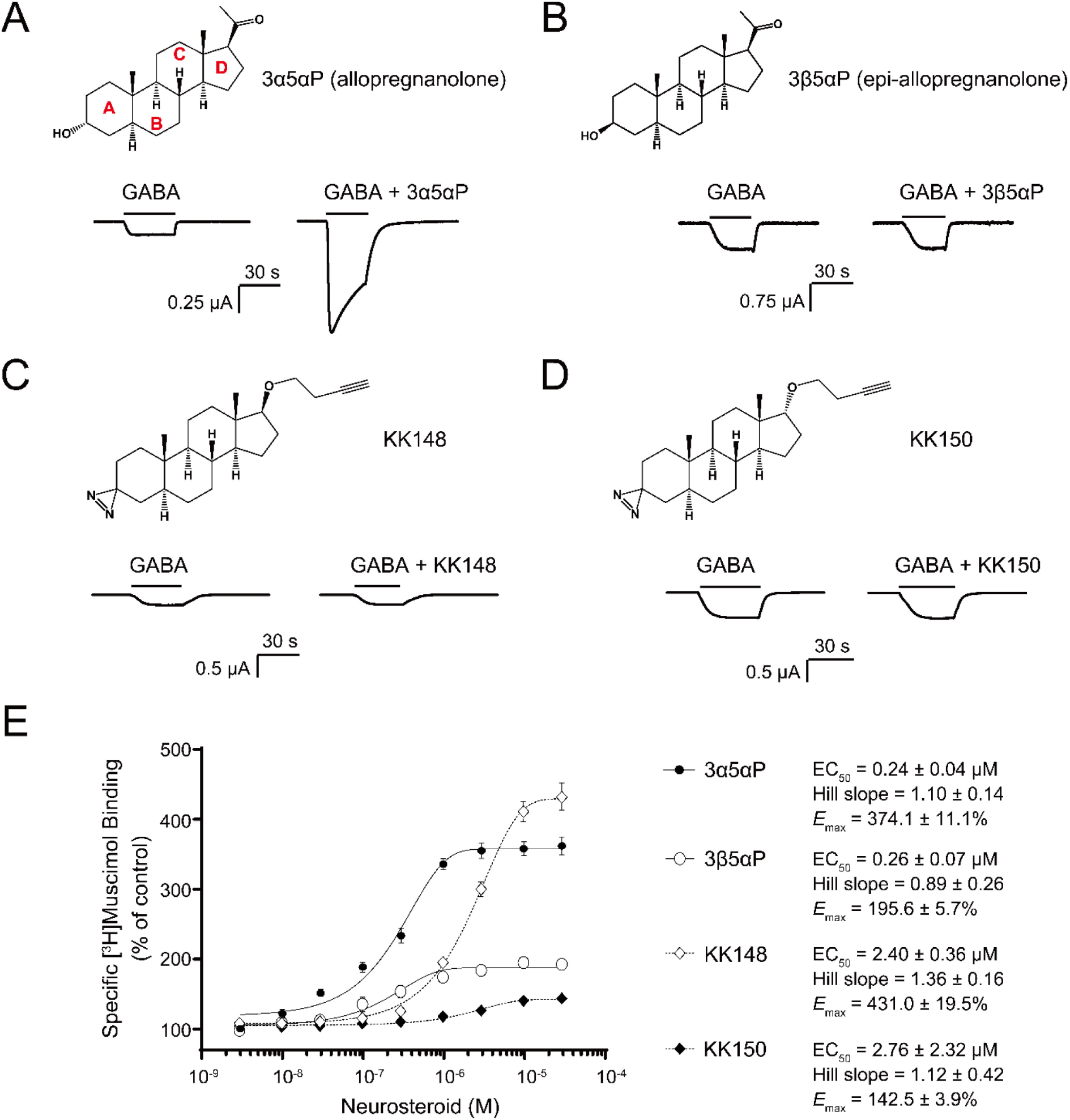
Distinct neurosteroid effects on potentiation of GABA_A_R currents and modulation of [^3^H]muscimol binding. (A) Structure of allopregnanolone (3α5αP) and sample current traces from α_1_β_3_ GABA_A_R activated by 0.3 μM GABA showing potentiation by 10 μM 3α5αP. (B), (C) and (D) Structures of epi-allopregnanolone (3β5αP), neurosteroid analogue photolabeling reagents KK148 and KK150, respectively, and sample current traces from α_1_β_3_ GABA_A_R activated by 0.3 μM GABA showing the absence of potentiation by 10 μM neurosteroids. (E) Concentration-response relationship for neurosteroid modulation of [^3^H]muscimol binding to α_1_β_3_ GABA_A_R. 3 nM–30 μM neurosteroids modulate [^3^H]muscimol (3 nM) binding in a concentration-dependent manner. Data points, EC_50_, Hill slope and maximal effect value [*E*_max_ (% of control): 100% means no effect] are presented as mean ± SEM (*n* = 6 for 3α5αP and KK148; *n* = 3 for 3β5αP and KK150). **Figure 1–figure supplement 1**: Neurosteroid modulation of muscimol binding to intact cells.

### State-specific actions of NS analogues

To determine why 3β5αP and KK148 enhance [^3^H]muscimol binding but do not potentiate α_1_β_3_ GABA_A_R currents, we first considered the possibility that 3β5αP- and KK148-induced enhancement of [^3^H]muscimol binding is a selective effect on intracellular GABA_A_Rs, since the radioligand binding assay was performed on total membrane homogenates, whereas the electrophysiological assays report only from cell surface channels. NS are known to have effects on intracellular GABA_A_Rs and have been shown to accelerate GABA_A_R trafficking (38-40). To test this possibility, we examined [^3^H]muscimol binding in intact cells (i.e. binding to receptors only in the plasma membrane) (41-43) compared to permeabilized cells (plasma membranes plus intracellular membranes). Notably, [^3^H]muscimol binding was two-fold greater in permeabilized cells than in intact cells, indicating a significant population of intracellular GABA_A_Rs. KK148 enhanced [^3^H]muscimol binding in intact cells as much or more than in permeabilized cells, indicating that this effect is not a result of selective NS actions on intracellular receptors (Figure 1–figure supplement 1).

A second possibility is that 3β5αP and KK148 selectively bind to and stabilize a desensitized state of the GABA_A_R. This would result in enhanced [^3^H]muscimol binding, since both open and desensitized receptors exhibit increased orthosteric ligand affinity in comparison to the resting state (11,18,37). To examine the effect of these NS analogues on desensitization, we maximally activated α_1_β_3_ GABA_A_R with 1 mM GABA and tested the effect of the NS on steady-state currents (44). KK148 and 3β5αP both decreased steady-state currents (Figures 2A and 2C), whereas KK150 did not (Figure 2B). To further delineate the electrophysiological effects of these compounds we focused on 3β5αP, since it is an endogenous NS and we had limited availability of KK148. 3β5αP preferentially inhibited steady-state rather than peak currents, indicating that inhibition is unlikely to be a rapidly-developing channel blocking effect. The inhibitory effect was also not observed in the α_1_(V256S)β_3_ TM2 pore-lining mutation, which was previously shown to remove the desensitizing effects of sulfated steroids (13,14) (Figure 2D). While both 3α5αP and 3β5αP enhance [^3^H]muscimol binding, the former predominantly results in receptor activation while the latter results in desensitization. Consistent with this, the α_1_(V256S)β_3_ mutation which abolishes NS-induced desensitization (13,14) eliminated [^3^H]muscimol binding enhancement by 3β5αP but not 3α5αP (Figure 3). We infer that 3α5αP increases [^3^H]muscimol binding by stabilizing an active state of the receptor, whereas 3β5αP increases [^3^H]muscimol binding by stabilizing a desensitized state of the receptor which is absent in the α_1_(V256S)β_3_ receptor. Collectively, these data indicate that 3β5αP and KK148 stabilize a desensitized state of the GABA_A_R, thus enhancing orthosteric ligand affinity.

**FIGURE 2:**
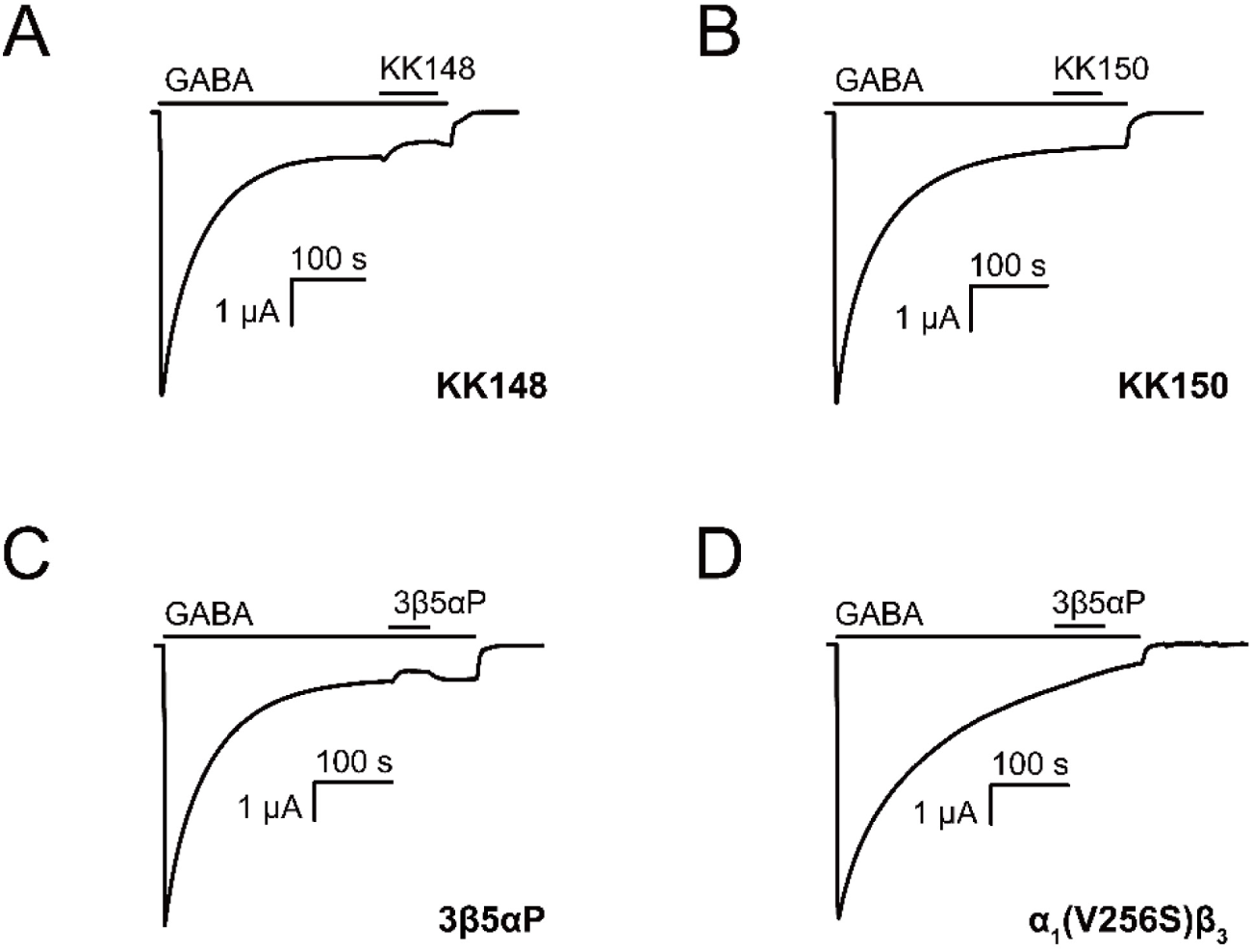
Neurosteroids promote steady-state desensitization of α_1_β_3_ GABA_A_Rs. Representative traces showing the effects of KK148, KK150 and epi-allopregnanolone (3β5αP) on maximal steady-state GABA-elicited currents. α_1_β_3_ GABA_A_Rs expressed in *Xenopus laevis* oocytes were activated with 1 mM GABA to maximally activate GABA_A_R current. (A-C) The effect of KK148 (10 μM), KK150 (10 μM) and 3β5αP (3 μM) on steady-state current. (D) The effect of 3β5αP (3 μM) on steady-state current in α_1_β_3_ GABA_A_Rs containing the α_1_V256S mutation, known to eliminate NS-induced desensitization. The results show that 3β5αP and KK148 reduce steady-state currents, consistent with enhanced desensitization, whereas KK150 does not. The effect of 3β5αP on steady-state currents is eliminated by the α_1_V256S mutation, consistent with 3β5αP enhancing desensitization rather than producing channel block.

**FIGURE 3:**
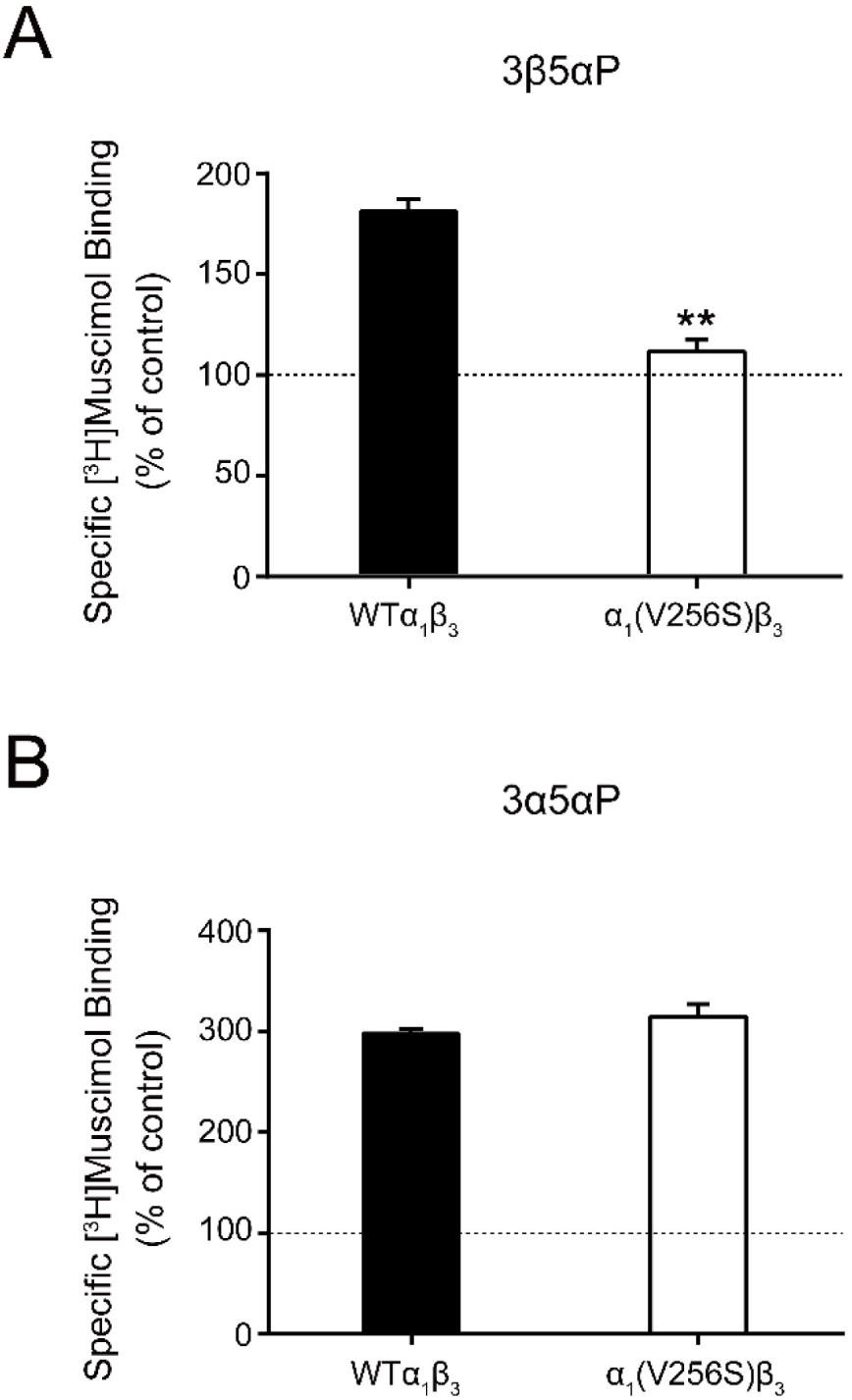
Effect of α_1_(V256S)β_3_ mutation on neurosteroid enhancement of [^3^H]muscimol binding. (A) Enhancement of specific [^3^H]muscimol (3 nM) binding to α_1_β_3_ GABA_A_R WT by 10 μM epi-allopregnanolone (3β5αP) is absent in α_1_(V256S)β_3_ GABA_A_R. (B) Enhancement of [^3^H]muscimol binding by 10 μM allopregnanolone (3α5αP) is unaffected by the α_1_V256S mutation. These data indicate that 3β5αP enhancement of orthosteric ligand binding requires receptor desensitization, whereas 3α5αP does not. (*n* = 6, ± SEM). \*\**P* < 0.01 vs. WT.

### Quantitative comparison of the effects of 3β5αP on [^3^H]muscimol binding and receptor desensitization

While there is qualitative agreement between the relative effects of the various NS analogues on orthosteric ligand binding and receptor desensitization, there is an apparent quantitative discrepancy in the magnitude of the effects. For example, 3β5αP enhances [^3^H]muscimol binding by two-fold (Figure 1E), whereas it reduces steady-state current by only ∼25% (Figure 2C). To address this difference, we first considered that the radioligand binding and electrophysiological assays are performed under different experimental conditions. The radioligand binding studies are performed using low [^3^H]muscimol concentrations to allow for sufficient dynamic range of ligand binding. In contrast the desensitization experiments are performed at high orthosteric ligand (GABA) concentration to achieve high peak open probability and steady-state receptor desensitization, thus minimizing the number of channels in the resting state. To address the quantitative differences in results from the two assays, we analyzed the electrophysiological data in the framework of the three-state Resting-Open-Desensitized model (44,45). We assumed that both the open and desensitized states had higher affinity for muscimol than the resting state, and that the affinities were similar and could be treated as equal. We then calculated the predicted occupancy of the high affinity states (P_open_ + P_desensitized_) using parameters derived from the functional responses, to compare to the observed changes in binding. The raw current amplitudes of peak and steady-state responses were converted to units of open probability as described previously in detail (46), and the probabilities of being in the open (P_open_) or desensitized (P_desensitized_) states were calculated for different experimental conditions (see Methods).

Application of 1 mM GABA elicited a current response that had a peak P_open_ of 0.71 ± 0.25 (*n* = 16). The P_open_ of the steady-state response was 0.121 ± 0.033 (*n* = 7), that was reduced to 0.077 ± 0.013 (*n* = 5) with 3 µM 3β5αP. Analysis of steady-state currents using the Resting-Open-Desensitized model indicates that the steady-state P_desensitized_ is 0.829 in the presence of GABA, and 0.892 in the presence of GABA + steroid. The relatively small increase in the sum of (P_open_ + P_desensitized_) (from 0.95 to 0.97) is due to the use of saturating GABA in these experiments. In the presence of lower concentrations of GABA, i.e., lower levels of activation, the predicted increase in the sum of (P_open_ + P_desensitized_) is greater. For example, for the condition where GABA elicits a peak response with P_open_ of 0.01, the predicted steady-state P_open_ is 0.0094 and the predicted P_desensitized_ 0.0643 (sum of the two equals 0.0737). In the presence of GABA + 3β5αP, the predicted steady-state P_open_ is 0.0090 and the predicted P_desensitized_ 0.1045 (sum of 0.1135), thus showing a 54% increase in the sum of (P_open_ + P_desensitized_). Incidentally, this example demonstrates the need to use high concentrations of GABA to observe a meaningful reduction in steady-state P_open_ in the presence of 3β5αP.

To more directly compare the data from the radioligand binding and electrophysiology experiments, we exposed oocytes containing α_1_β_3_ GABA_A_Rs to 20 nM muscimol and recorded currents before and after co-application of 3 μM 3β5αP. The percent reduction in steady-state current following 3β5αP exposure was measured and used to estimate the relative probabilities of closed, open and desensitized receptors. The application of 20 nM muscimol elicited a peak response with P_open_ of 0.012 ± 0.004 (*n* = 6). The steady-state P_open_ was 0.011 ± 0.004. In the same cells, subsequent exposure to 3 µM 3β5αP reduced the steady-state P_open_ to 0.009 ± 0.004. The calculated steady-state P_desensitized_ was 0.1001 in the presence of muscimol, and 0.2168 in the presence of muscimol + 3β5αP. Thus, there is a predicted two-fold increase in the sum of (P_open_ + P_desensitized_) when the steroid is combined with muscimol. This is consistent with the doubling of muscimol binding caused by 3β5αP in the [^3^H]muscimol binding experiments (Figure 1E). Overall, the data indicate that relatively small changes in steady-state current can be associated with relatively large changes in the occupancy of high-affinity states.

### Binding site selectivity for NS analogues

To determine whether KK148 and 3β5αP stabilize the desensitized conformation of the GABA_A_R by selectively binding to one or more of the identified NS binding sites on the GABA_A_R (11), we first determined which of the identified NS sites they bind. We have previously shown that the 3α5αP-analogue photolabeling reagent, KK200 labels the β_3_(+)–α_1_(-) intersubunit (β_3_G308) and α_1_ intrasubunit (α_1_N408) sites on α_1_β_3_ GABA_A_Rs (Figure 4A), and that photolabeling can be prevented by a ten-fold excess of 3α5αP (11). As a first step to determine the binding sites for 3β5αP, KK148 or KK150, we examined whether a ten-fold excess of these compounds (30 μM) prevented KK200 (3 μM) photolabeling of either binding site. Photolabeling was performed on membranes from HEK293 cells transfected with epitope-tagged α_1His-FLAG_β_3_ receptors, mimicking the conditions used in the [^3^H]muscimol binding assays and photolabeled residues were identified and labeling efficiency was determined using middle-down mass spectrometry (11). KK148, KK150, 3α5αP and 3β5αP all prevented KK200 photolabeling of α_1_N408 in the α_1_ intrasubunit site (Figure 4B), consistent with their binding to this site. In contrast, KK148, KK150 and 3α5αP but not 3β5αP prevented labeling of β_3_G308 in the intersubunit site (Figure 4C), indicating that 3β5αP does not bind to the intersubunit site.

**FIGURE 4:**
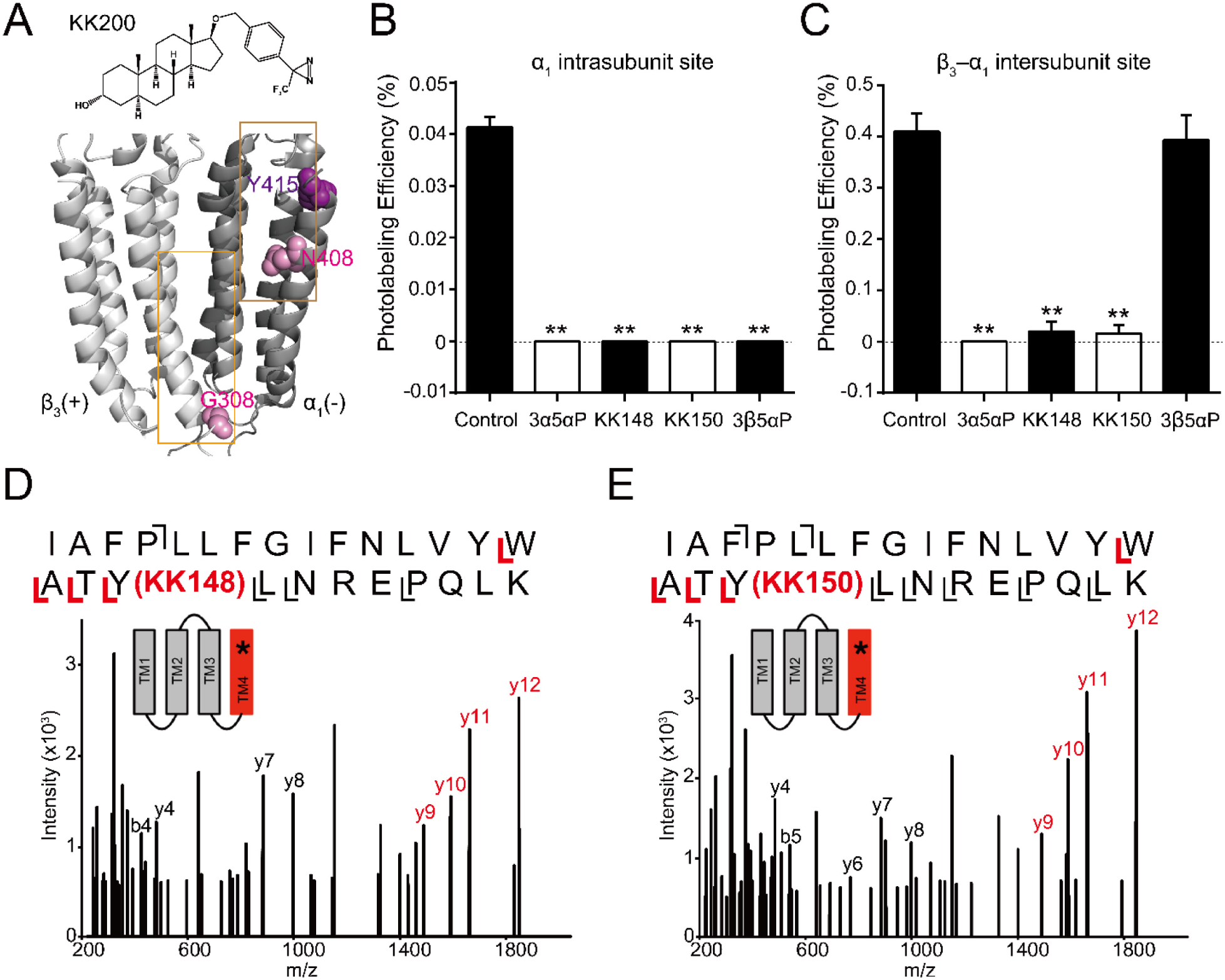
Competitive prevention of neurosteroid photolabeling at an intersubunit and intrasubunit site. (A) Structures of the neurosteroid photolabeling reagent KK200 and the α_1_β_3_ GABA_A_R-TMDs highlighting the residues G308 in the β_3_(+)–α_1_(-) intersubunit site and N408 in the α_1_ intrasubunit site previously identified by KK200 photolabeling in pink. Shown in purple is Y415 in the α_1_ intrasubunit site, which is photolabeled by KK148 and KK150. Adjacent β_3_(+) and α_1_(-) subunits are shown and the channel pore is behind the subunits. (B) Photolabeling efficiency of α_1_ subunit TM4 (α_1_ intrasubunit site) in α_1_β_3_ GABA_A_R by 3 μM KK200 in the absence or presence of 30 μM allopregnanolone (3α5αP), KK148, KK150, and epi-allopregnanolone (3β5αP) (*n* = 3, ± SEM). \*\**P* < 0.01 vs. control. (C) Same as (B) for β_3_ subunit TM3 [β_3_(+)–α_1_(-) intersubunit site, *n* = 3, ± SEM]. (D) HCD fragmentation spectrum of the α_1_ subunit TM4 tryptic peptide photolabeled by 30 μM KK148. Red and black indicate fragment ions that do or do not contain KK148, respectively. The schematic highlight in red identifies the TMD being analyzed and the asterisk denotes the approximate location of KK148. (E) Same as (D) photolabeled by 30 μM KK150. **Figure 4–figure supplement 1**: Extracted ion chromatograms of labeled and unlabeled β_3_ subunit TM4 peptides. **Figure 4–figure supplement 2**: Fragmentation spectrum of unlabeled α_1_ subunit TM4 peptide.

The KK148- and KK150-photolabeled samples were also analyzed to directly identify the sites of adduction. In both the KK148- and KK150-labeled samples, photolabeled peptides were identified from the TM4 helices of both the α_1_ and β_3_ subunits. The labeled peptides had longer chromatographic elution times than the corresponding unlabeled peptides and corresponded with high mass accuracy (< 20 ppm) to the predicted mass of the unlabeled peptides plus the add weight minus N_2_ of KK148 or KK150 (Figure 4– figure supplement 1). Product ion (MS2) spectra of the KK148- and KK150-labeled peptides from the α_1_ subunit identified the labeled residue as Y415 for both KK148 and KK150 (Figures 4D-E, Figure 4–figure supplement 2); Y415 is the same residue labeled by KK123 at the α_1_ intrasubunit site (11). The KK148 and KK150 labeled peptides in TM4 of the β_3_ subunit and corresponding unlabeled peptide were identified by fragmentation spectra as β_3_TM4 I426-N445. These data support labeling of the β_3_ intrasubunit site by KK148 and KK150. Fragmentation spectra of the peptide-sterol adducts were not adequate to determine the precise labeled residue because of low photolabeling efficiency (0.13% for KK148; 0.19% for KK150, Figure 4–figure supplement 1). No photolabeled peptides were identified in the β_3_(+)–α_1_(-) intersubunit site. This is likely because KK148 and KK150, similar to KK123, utilize an aliphatic diazirine that preferentially labels nucleophilic residues (28,34,47); such residues are not present in the intersubunit site.

We have also shown that KK123 labeling of the α_1_ intrasubunit (α_1_Y415) and β_3_ intrasubunit (β_3_Y442) sites (Figure 5A) can be prevented by a ten-fold excess of 3α5αP (11). We thus examined whether 3β5αP (30 μM) inhibited photolabeling by KK123 (3 μM). 3β5αP completely inhibited KK123 photolabeling at both intrasubunit sites (Figures 5B-C). Collectively the data show that KK148, KK150 and 3α5αP bind to all three of the identified NS binding sites. In contrast, 3β5αP selectively binds to the two intrasubunit binding sites, but not to the canonical β_3_(+)–α_1_(-) intersubunit site.

**FIGURE 5:**
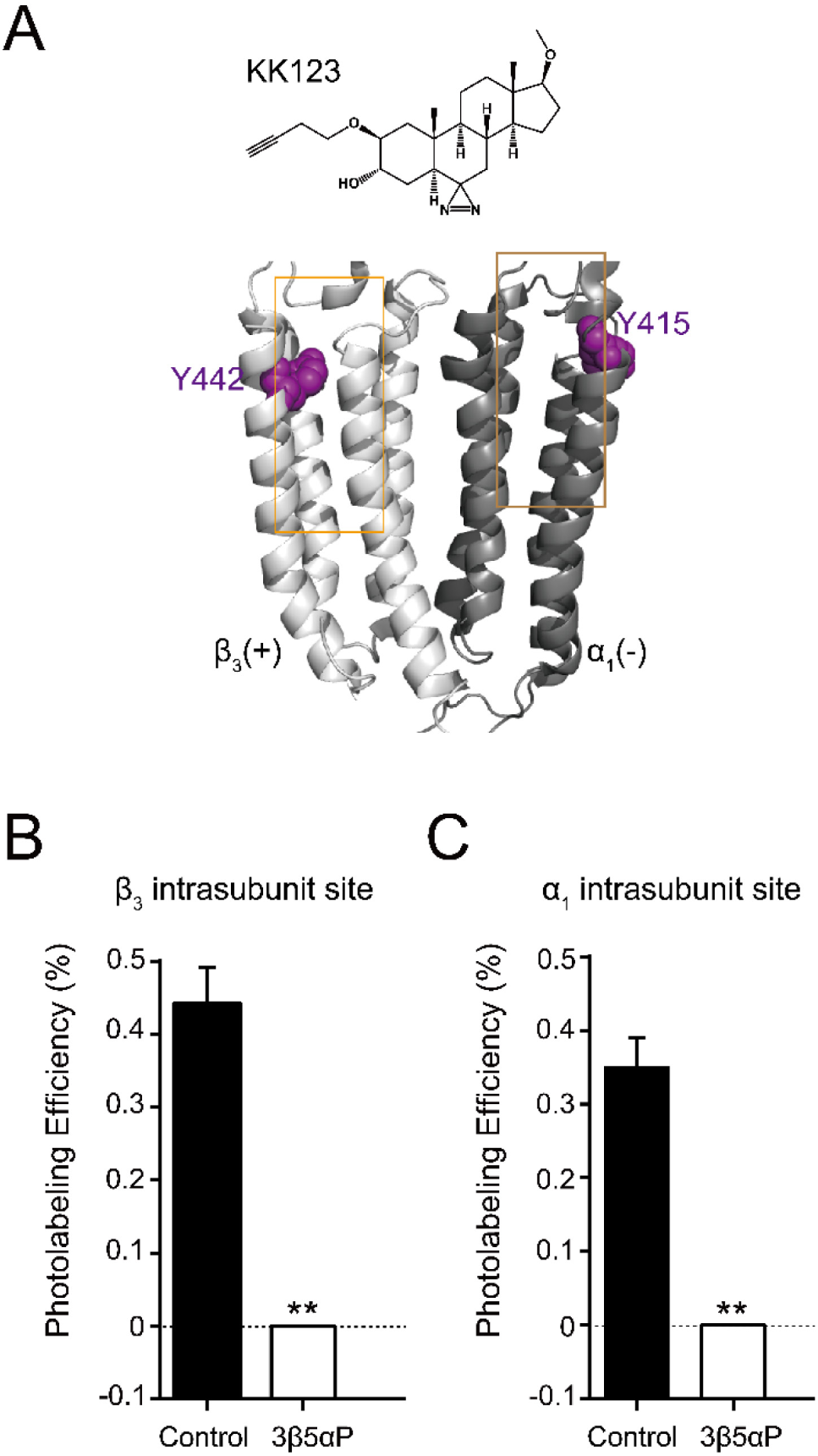
Epi-allopregnanolone prevents neurosteroid photolabeling at the α_1_ and β_3_ intrasubunit sites. (A) Structures of the neurosteroid photolabeling reagent KK123 and the α_1_β_3_ GABA_A_R-TMDs highlighting the residues Y442 in the β_3_ intrasubunit site and Y415 in the α_1_ intrasubunit site previously identified by KK123 photolabeling in purple. Adjacent β_3_(+) and α_1_(-) subunits are shown and the channel pore is behind the subunits. (B) Photolabeling efficiency of β_3_ subunit TM4 (β_3_ intrasubunit site) in α_1_β_3_ GABA_A_R by 3 μM KK123 in the absence or presence of 30 μM epi-allopregnanolone (3β5αP) (*n* = 3, ± SEM). \*\**P* < 0.01 vs. control. (C) Same as (B) for α_1_ subunit TM4 (α_1_ intrasubunit site, *n* = 3, ± SEM).

### Orthosteric ligand binding enhancement by NS analogues is mediated by distinct sites

To determine which of the previously identified binding sites contributes to NS enhancement of [^3^H]muscimol binding, we performed site-directed mutagenesis of the NS binding sites previously determined by photolabeling (Figure 6A) (11). Specifically, α_1_(Q242L)β_3_ targets the β_3_(+)–α_1_(-) intersubunit site, α_1_(N408A/Y411F)β_3_ and α_1_(V227W)β_3_ the α_1_ intrasubunit site, and α_1_β_3_(Y284F) the β_3_ intrasubunit site. Mutations in the β_3_(+)–α_1_(-) intersubunit and α_1_ intrasubunit sites decreased 3α5αP enhancement of [^3^H]muscimol binding by ∼80%, while mutation of the β_3_ intrasubunit site led to a small decrease (Figure 6B, Table 1). The residual enhancement of [^3^H]muscimol binding observed in receptors with mutations in the intersubunit or α_1_ intrasubunit site occurs at ten-fold higher concentrations of 3α5αP than WT and receptors with mutations in the β_3_ intrasubunit site (Table 1), suggesting that 3α5αP binds to the β_3_ intrasubunit site with lower affinity. In contrast, mutations in the α_1_ and β_3_ intrasubunit sites, but not the intersubunit site decreased the enhancement of [^3^H]muscimol binding by 3β5αP and KK148 (Figures 6C-D, Table 1). To confirm that the effect of these mutations on NS effect are steroid-specific, we also tested their effect on etomidate, which enhances [^3^H]muscimol binding in α_1_β_3_ GABA_A_Rs and acts through a binding site distinct from NS (48,49). The mutations targeting NS binding sites resulted in modest decreases in [^3^H]muscimol binding enhancement by etomidate; however, the α_1_β_3_(M286W) mutation which abolishes etomidate potentiation and activation of GABA_A_Rs (50,51), also abolished [^3^H]muscimol binding enhancement (Figure 6E).

**TABLE 1:**
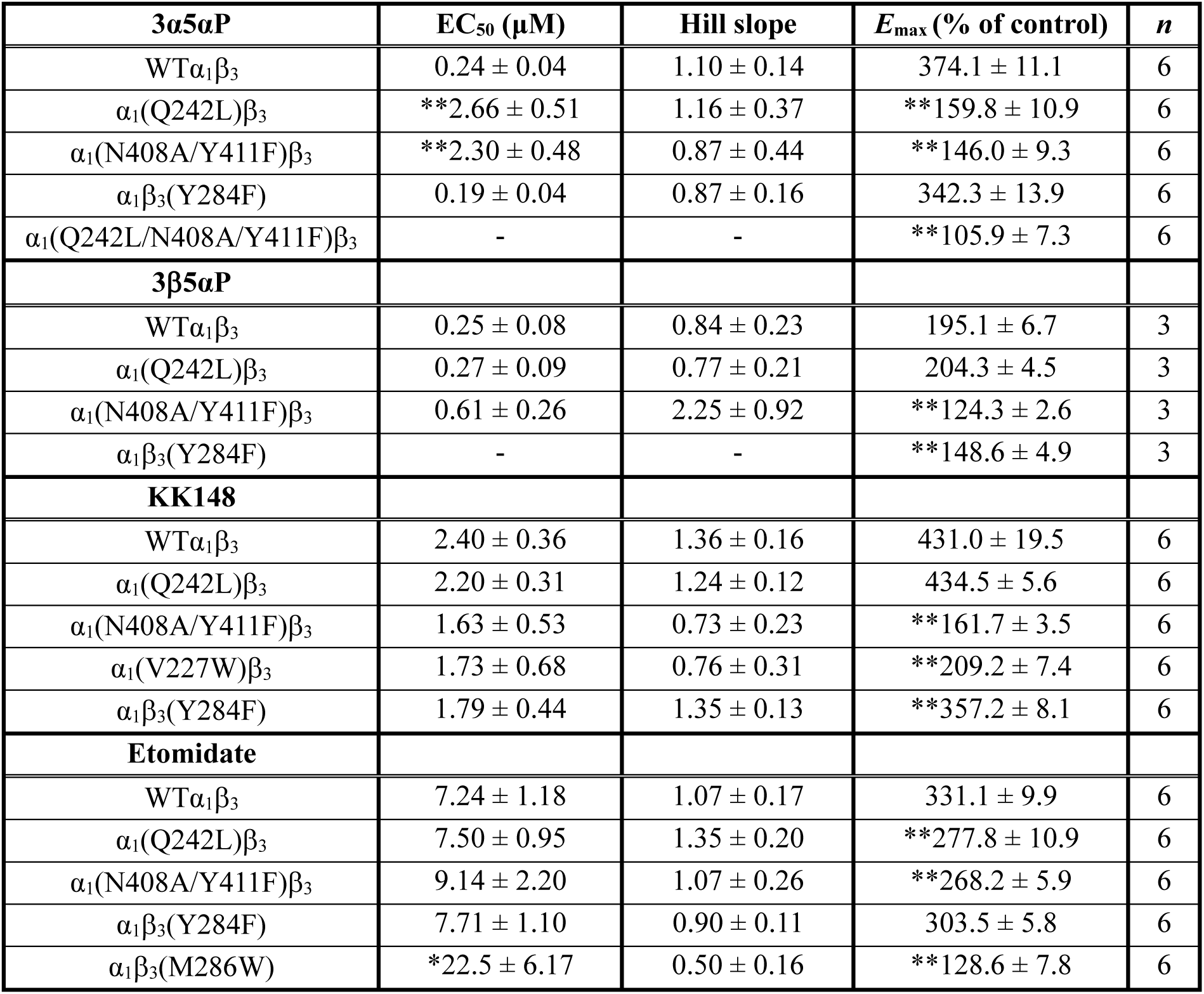
Effects of mutations on neurosteroid modulation of [^3^H]muscimol binding. EC_50_, Hill slope and maximal effect values [*E*_max_ (% of control): 100% means no effect] for the concentration-response curves in Figures 6B-E. Statistical differences are analyzed using one-way ANOVA with *post hoc* Bonferroni correction (\**P* < 0.05 vs. WT; \*\**P* < 0.01 vs. WT). Data are presented as mean ± SEM.

**FIGURE 6:**
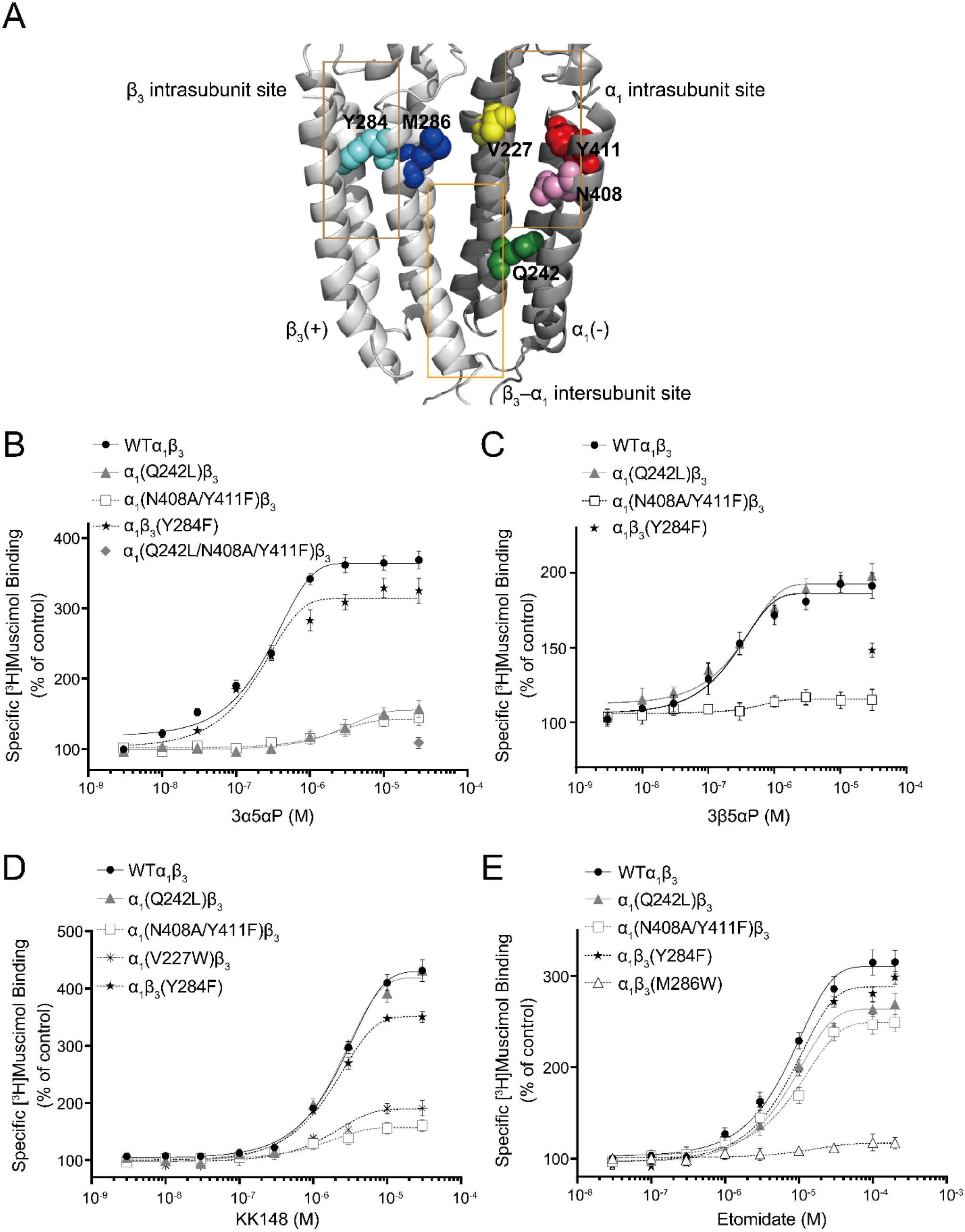
Effect of mutations in α_1_β_3_ GABA_A_R on neurosteroid modulation of [^3^H]muscimol binding. (A) Structure of the α_1_β_3_ GABA_A_R-TMD highlighting the residues where mutations were made in putative binding sites for neurosteroids (Q242-green for β_3_–α_1_ intersubunit site; V227-yellow, N408-pink and Y411-red for α_1_ intrasubunit site; Y284-cyan for β_3_ intrasubunit site) and M286-blue for etomidate. Adjacent β_3_(+) and α_1_(-) subunits are shown and the channel pore is behind the subunits. (B) Concentration-response relationship for the effect of 3 nM–30 μM allopregnanolone (3α5αP) on [^3^H]muscimol (3 nM) binding to α_1_β_3_ GABA_A_R WT and indicated mutants. Data points represent mean ± SEM (*n* = 6). (C), (D) and (E) Same as (B) for 3 nM–30 μM epi-allopregnanolone (3β5αP) (*n* = 3), KK148 (*n* = 6) and 30 nM–200 μM etomidate (*n* = 6), respectively. The data for WT in panels 6B and 6D is a replot of the same data shown in Figure 1E. **Figure 6–figure supplement 1**: Neurosteroid effect on [^3^H]muscimol binding isotherms in α_1_β_3_ WT and α_1_(N408A/Y411F)β_3_ GABA_A_Rs. **Figure 6–figure supplement 2**: Properties of [^3^H]muscimol saturation binding curves in the α_1_β_3_ GABA_A_R WT and α_1_(N408A/Y411F)β_3_ mutant.

We did not test the effects of mutations on KK150 action because it minimally enhances [^3^H]muscimol binding. However, KK150 binds to all three of the identified NS binding sites, and may thus be a weak partial agonist or antagonist at the sites mediating NS enhancement of [^3^H]muscimol binding. Consistent with this prediction, KK150 inhibited enhancement of [^3^H]muscimol binding by 3α5αP and KK148 (Figure 7).

**FIGURE 7:**
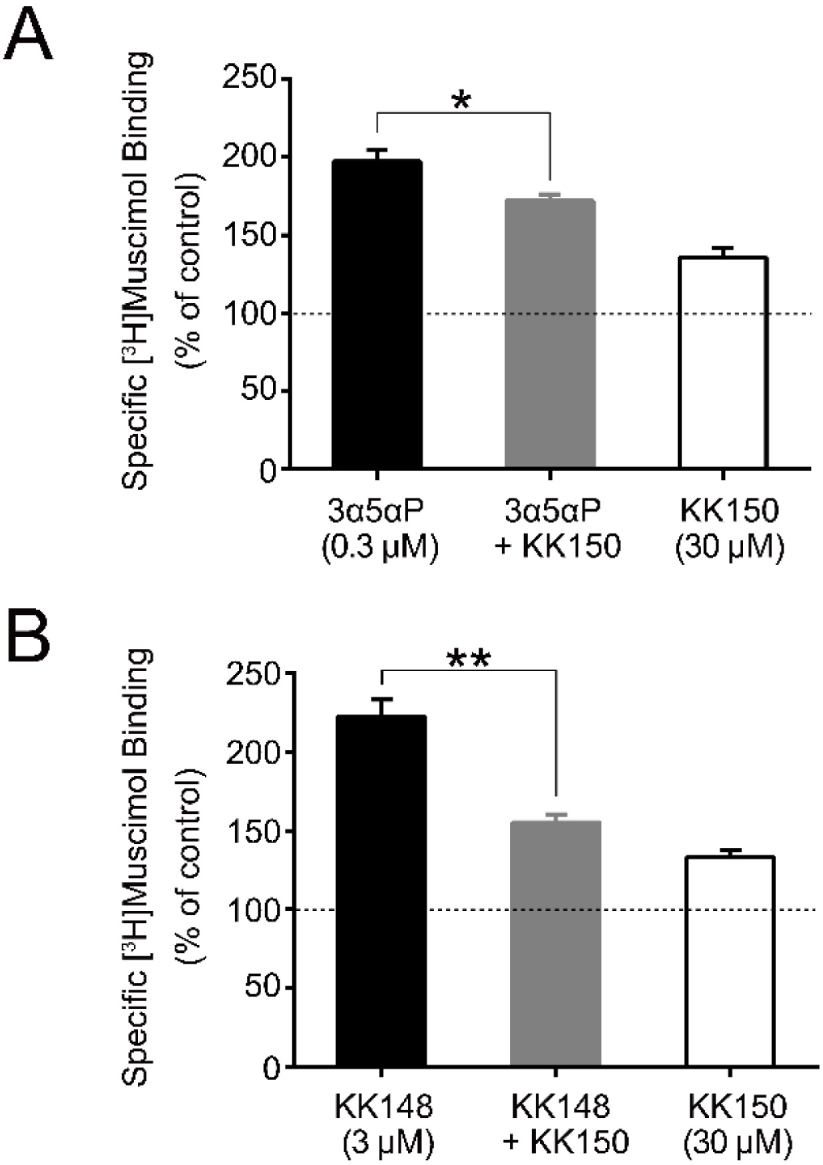
KK150 prevents neurosteroid-induced muscimol binding enhancement. (A) Enhancement of specific [^3^H]muscimol (3 nM) binding to α_1_β_3_ GABA_A_R by 0.3 μM allopregnanolone (3α5αP) in the absence (black bar) or presence (grey bar) of 30 μM KK150 and KK150 alone (white bar) (*n* = 8, ± SEM). \**P* < 0.05 vs. 0.3 μM 3α5αP alone. (B) Same as (A) for 3 μM KK148 (*n* = 8, ± SEM). \*\**P* < 0.01 vs. 3 μM KK148 alone.

Collectively, these results show that multiple NS binding sites contribute to enhancement of [^3^H]muscimol affinity and that potentiating NS (3α5αP) and non-potentiating NS (3β5αP, KK148 and KK150) have both common and distinct sites of action. Specifically, 3α5αP enhances [^3^H]muscimol binding through all three sites but predominantly through the intersubunit and α_1_ intrasubunit sites, which we have previously shown mediate PAM-NS potentiation (11). In contrast, 3β5αP and KK148 enhance [^3^H]muscimol binding exclusively through the α_1_ and β_3_ intrasubunit sites. KK150 antagonizes the effects of KK148 on [^3^H]muscimol binding, presumably via the intrasubunit sites, and antagonizes the effects of 3α5αP, possibly via all three sites. These data indicate that NS binding to both the intersubunit and intrasubunit sites contributes to 3α5αP enhancement of [^3^H]muscimol binding, but that only the intrasubunit binding sites contribute to the effects of 3β5αP and KK148.

It is important to note that the [^3^H]muscimol binding curves in Figure 6 are normalized to control. The raw data show that membranes containing WT receptors have 10-20-fold higher [^3^H]muscimol binding (B_max_) than membranes containing α_1_(N408A/Y411F)β_3_ receptors (Figure 6–figure supplement 1A). The lower total [^3^H]muscimol binding observed in α_1_(N408A/Y411F)β_3_ membranes is likely a consequence of decreased receptor expression. To assure that differences in NS effect between WT and α_1_(N408A/Y411F)β_3_ are not due to different muscimol affinities, we examined [^3^H]muscimol binding at a full range of concentrations. The α_1_(N408A/Y411F)β_3_ mutations did not have a significant effect on [^3^H]muscimol affinity (Figure 6–figure supplement 1B), but eliminated the modulatory effects of NS (3α5αP and KK148) on [^3^H]muscimol affinity (Figure 6–figure supplement 1C-D and Figure 6–figure supplement 2). To assure that the effect of α_1_(N408A/Y411F)β_3_ was specific to NS, we also examined the effect of etomidate (a non-steroidal GABA_A_R PAM) on muscimol affinity. Etomidate enhanced [^3^H]muscimol affinity in both the WT and α_1_(N408A/Y411F)β_3_ receptors, indicating that the effect of these mutations are specific to NS action (Figure 6–figure supplement 1C-D).

### 3β5αP increases desensitization through binding to α_1_ and β_3_ intrasubunit sites

To further explore the relationship between desensitization and enhancement of [^3^H]muscimol binding, we examined the consequences of mutations to these sites on physiological measurements of desensitization induced by NS. Again, these experiments were performed with 3β5αP because it is the endogenous 3β-OH NS and because of limited quantities of KK148. Desensitization was quantified by defining the baseline steady-state current at 1 mM GABA as 100% and measuring percent reduction of the steady-state current elicited by a NS (Figure 8A). 3β5αP reduced the steady-state current (i.e. enhanced desensitization) by 23.0 ± 2.8% (% of desensitization: mean ± SEM, *n* = 5, Figure 8A). Mutations in the α_1_ and β_3_ intrasubunit sites [i.e. α_1_(N408A/Y411F)β_3_ and α_1_β_3_(Y284F), respectively] prevented 3β5αP-enhanced desensitization by ∼67% (Figure 8B), whereas mutation in the β(+)–α(-) intersubunit site [α_1_(Q242L)β_3_] was without effect (Figure 8C). Receptors with mutations in both the α_1_ and β_3_ intrasubunit sites [α_1_(N408A/Y411F)β_3_(Y284F)] showed less NS-enhancement of desensitization than receptors with mutations in either of the intrasubunit sites alone, indicating that both intrasubunit sites contribute to the desensitizing effect (Figure 8C). Whereas the desensitizing effect of 3β5αP is completely eliminated by the V2’S mutation α_1_(V256S)β_3_, it is not completely eliminated by combined mutations of all three binding sites [α_1_(Q242L/N408A/Y411F)β_3_(Y284F)] (Figure 8C). This suggests either that the effects of the mutations are incomplete or there are additional unidentified NS binding sites contributing to desensitization. Since mutations of the α_1_ and β_3_ intrasubunit sites also disrupt 3β5αP-enhancement of [^3^H]muscimol binding (Figure 6C), we conclude that 3β5αP binding to these intrasubunit sites stabilizes the desensitized state of the GABA_A_R and thus enhances [^3^H]muscimol binding. Furthermore, KK148 increased GABA_A_R desensitization (% of desensitization = 27.2 ± 6.0: mean ± SEM, *n* = 3, Figure 2A) and the α_1_(V256S)β_3_ mutation abolished the effect (% of desensitization = 0, *n* = 1). These observations support the idea that binding of certain NS analogues to α_1_ and β_3_ intrasubunit sites, increases GABA_A_R desensitization. In contrast, KK150 showed a very small effect on desensitization (% of desensitization = 2.1 ± 0.7: mean ± SEM, *n* = 5, Figure 2B), consistent with the small increase in [^3^H]muscimol binding by KK150 (Figure 1E).

**FIGURE 8:**
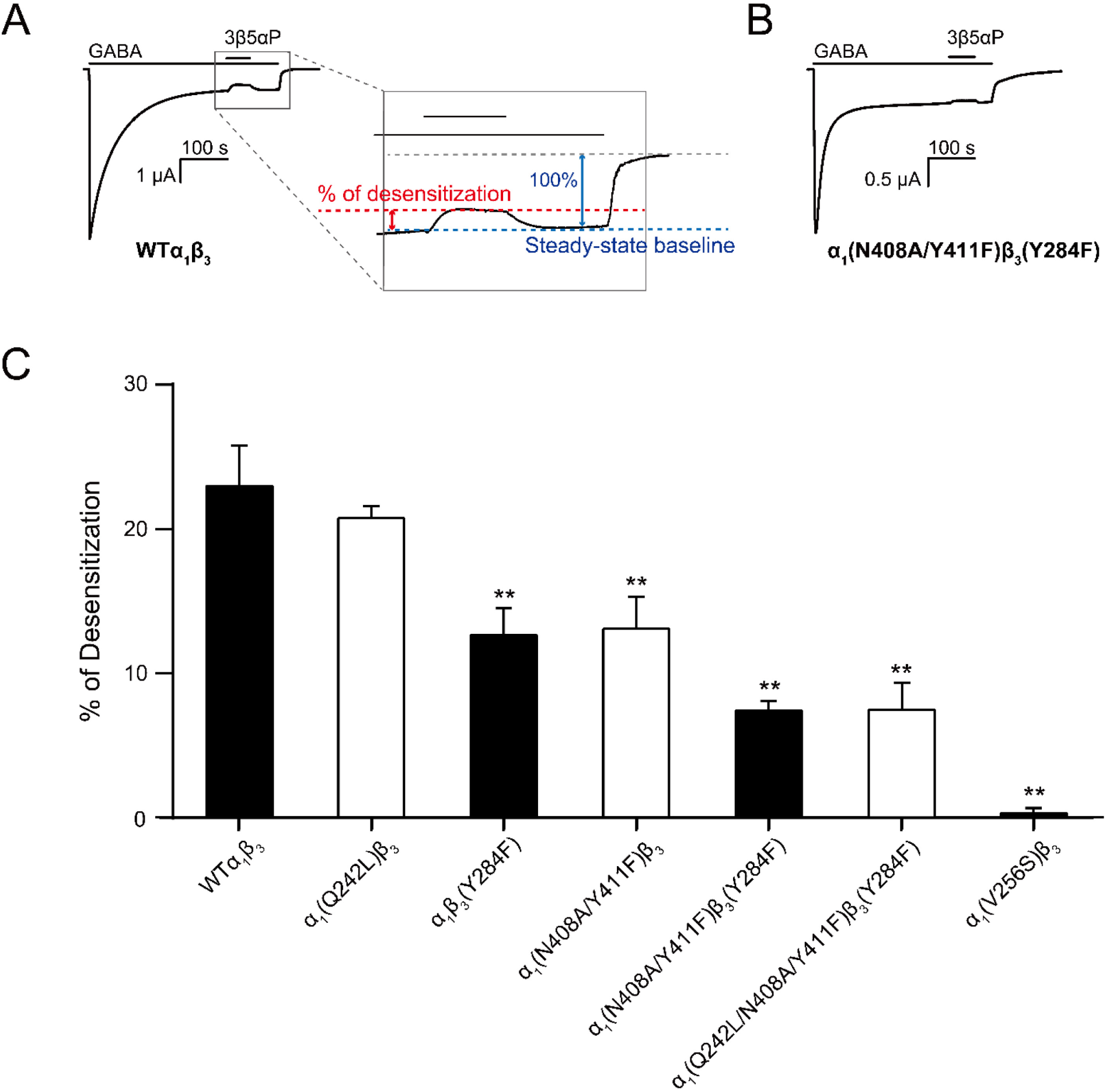
Mutations in intrasubunit sites prevent desensitization by epi-allopregnanolone. (A) Sample current trace showing the effect of 3 μM epi-allopregnanolone (3β5αP) on α_1_β_3_ GABA_A_R activated by 1 mM GABA. A zoomed-in box shows neurosteroid-induced desensitization of the steady-state GABA current. (B) Same as (A) for α_1_(N408A/Y411F)β_3_(Y284F) GABA_A_R. (C) Percent desensitization of the steady-state α_1_β_3_ GABA_A_R currents (WT and mutants) by 3 μM 3β5αP during continuous application of 1 mM GABA. (*n* = 5, ± SEM). \*\**P* < 0.01 vs. WT.

### The effects of 3α5αP binding to intrasubunit sites on desensitization

3α5αP binds to all three of the NS binding sites on α_1_β_3_ GABA_A_R, and mutations in all three sites reduce 3α5αP enhancement of [^3^H]muscimol binding (Figure 6B). This suggests the possibility that activation by 3α5αP (mediated primarily by the β_3_(+)–α_1_(-) intersubunit site) masks a desensitizing effect mediated through the β_3_ and/or α_1_ intrasubunit binding sites. To determine whether intrasubunit binding sites mediate increased desensitization by 3α5αP, we examined the effect of 3α5αP on steady-state currents in receptors with mutations in the α_1_ or β_3_ intrasubunit site. In these experiments, P_open_ was maximized by activating receptors with 1 mM GABA co-applied with 40 μM pentobarbital (PB) (52) prior to application of 3α5αP; this was necessary because several of the mutated receptors had a P_open_ << 1.0 in response to 1 mM GABA alone. Mutations in the intrasubunit sites were prepared with a background α_1_(Q242L)β_3_ mutation to remove 3α5αP activation (11,28,29,53) and focus on the effects of 3α5αP on the equilibrium between the open and desensitized states.

3α5αP produced a small reduction in steady-state current in α_1_(Q242L)β_3_ receptors with mutations in neither of the intrasubunit sites (Figure 9A). This inhibitory effect was eliminated by α_1_(V256S)β_3_, indicating that it was due to receptor desensitization (Figure 9D). In receptors with combined mutations in the intersubunit and α_1_ intrasubunit sites [i.e. α_1_(Q242L/N408A/Y411F)β_3_], 3α5αP significantly inhibited the steady-state current (Figure 9B), an effect that was markedly reduced by mutations in the β_3_ intrasubunit site [α_1_(Q242L)β_3_(Y284F)] (Figure 9C). These data suggest that 3α5αP exerts a desensitizing effect by binding to the β_3_ intrasubunit site and that 3α5αP binding to the α_1_ intrasubunit site does not promote desensitization (Figure 9D). Notably, 3α5αP exerted only a modest inhibitory effect in α_1_(Q242L)β_3_ receptors in which occupancy of the β_3_ intrasubunit site should promote inhibition. This may be due to a counterbalancing action at the α_1_ intrasubunit site, where 3α5αP binding contributes more to receptor activation as demonstrated by our previous observation that mutations in the α_1_ intrasubunit site significantly reduce 3α5αP potentiation of GABA-elicited currents (11). These results suggest that in addition to activation, 3α5αP enhances receptor desensitization. Enhanced desensitization by the PAM-NS 3α5αP (54) and 3α5α-THDOC (55,56) has been observed in prior studies supporting the current finding with 3α5αP.

**FIGURE 9:**
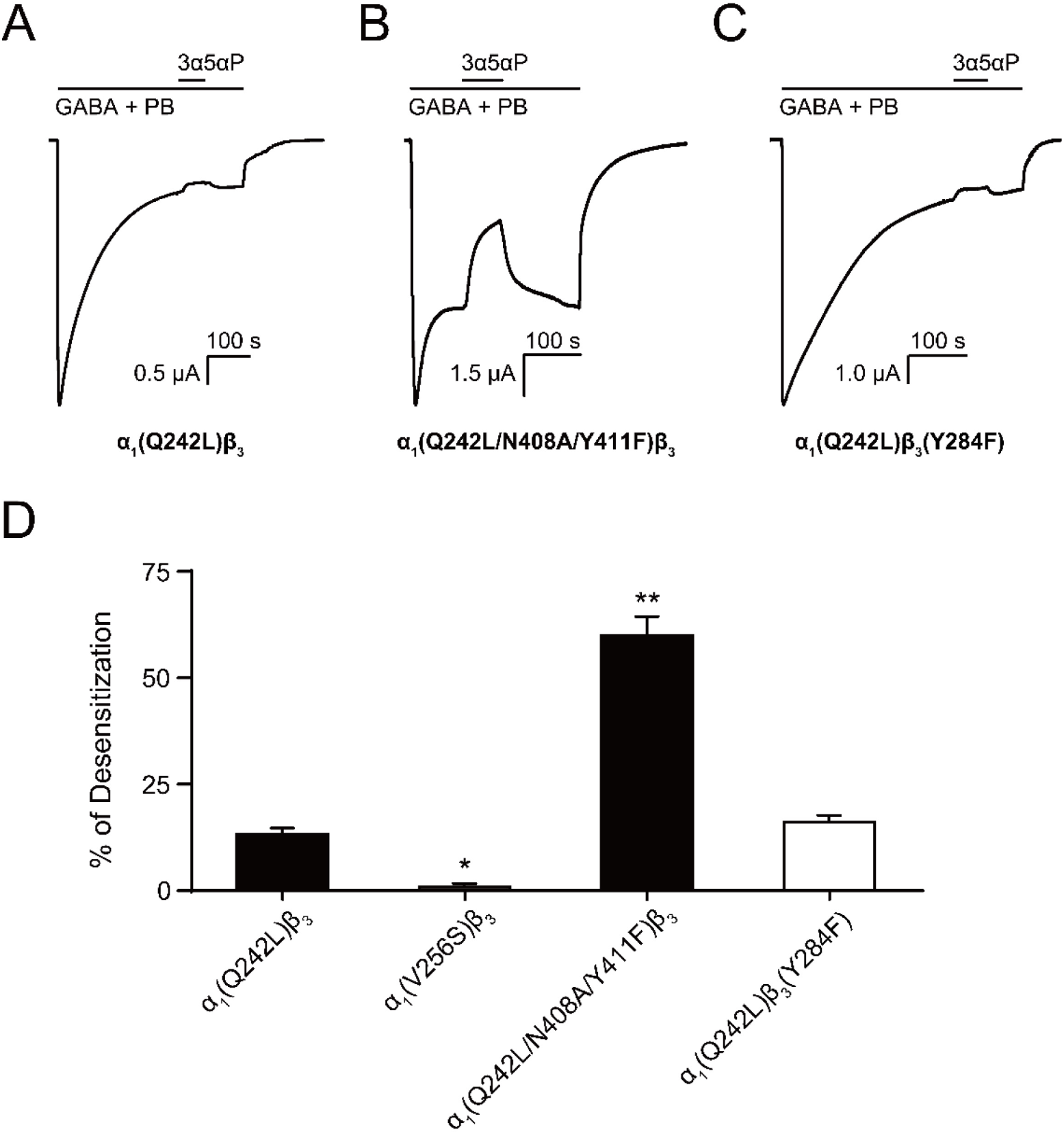
Allopregnanolone desensitizes GABA_A_R currents via binding to the β_3_ intrasubunit site. (A) Sample current trace showing the effect of 3 μM allopregnanolone (3α5αP) on α_1_(Q242L)β_3_ GABA_A_R activated by 1 mM GABA co-applied with 40 μM pentobarbital (PB). (B), (C) Same as (A) for α_1_(Q242L/N408A/Y411F)β_3_ GABA_A_R and α_1_(Q242L)β_3_(Y284F) GABA_A_R, respectively. (D) Percent desensitization of the steady-state currents elicited by 1 mM GABA with 40 μM PB in α_1_β_3_ GABA_A_R with specified mutations [*n* = 4 for α_1_(V256S)β_3_; *n* = 5 for others, ± SEM]. \**P* < 0.05; \*\**P* < 0.01 vs. α_1_(Q242L)β_3_, respectively.

## DISCUSSION

In this study we examined how site-specific binding to the three identified NS sites on α_1_β_3_ GABA_A_R (11) contributes to the PAM vs. NAM activity of epimeric 3-OH NS. We found that the PAM-NS 3α5αP, but not the NAM-NS 3β5αP, binds to the canonical β_3_(+)–α_1_(-) intersubunit site that mediates receptor potentiation, explaining the absence of 3β5αP PAM activity. In contrast, 3β5αP binds to intrasubunit sites in the α_1_ and β_3_ subunits, promoting receptor desensitization. Binding to the intrasubunit sites provides a mechanistic explanation for the NAM effects of 3β5αP (14). 3α5αP also binds to the β_3_ intrasubunit site explaining the previously described desensitizing effect of the PAM-NS 3α5αP (54) and 3α5α-THDOC (55,56). Two synthetic NS with diazirine moieties at C3 (KK148 and KK150) were used to identify NS binding sites and shown to bind to the intersubunit as well as both intrasubunit sites. Neither of these ligands potentiated agonist-activated GABA_A_R currents, reinforcing the importance of the 3α-OH group and its interaction with α_1_Q242 in PAM actions. KK148 is an efficacious desensitizing agent, acting through the α_1_ and β_3_ intrasubunit NS binding sites. KK150, the 17α-epimer of KK148, binds to all three NS binding sites, but neither activates nor desensitizes GABA_A_Rs, suggesting a potential chemical scaffold for a general NS antagonist. Collectively, these data show that differential occupancy of and efficacy at three discrete NS binding sites determines whether a NS ligand has PAM, NAM, or potentially NS antagonist activity on GABA_A_Rs.

The observation that 3β5αP and KK148 enhance orthosteric ligand binding but do not potentiate GABA-elicited currents first suggested that these NAM-NS selectively stabilize a desensitized conformation of the receptor (n.b. there may be multiple desensitized conformations of the receptor, possibly including NS-specific desensitized states, and our data does not address the specific conformations stabilized by NAM-NS. Strictly, our data indicate that the steroids stabilize a non-conducting state that has high affinity to the orthosteric agonist muscimol.) Chang and colleagues have shown that orthosteric ligand affinity (muscimol or GABA) is greater in desensitized and activated (open) GABA_A_Rs than in resting (closed) receptors, with estimated GABA K_D_ values of 78.5 μM, 120 nM and 40 nM for the resting, activated and desensitized α_1_β_2_γ_2_ receptors respectively (37). Our experimental and modeling data demonstrate that NAM-NS such as 3β5αP or KK148 enhance orthosteric ligand binding by increasing the population of receptors in a desensitized state. It is, however, unclear if 3β5αP or KK148 can promote transition of resting receptors directly to a desensitized state, thus bypassing channel opening. We propose that in the presence of low concentrations of orthosteric agonists (as in the [^3^H]muscimol binding assays), there is a slow shift of receptors from resting through activated to a desensitized state with minimal change in the population of receptors in the activated state. The slow time course of accumulation of desensitized receptors is illustrated by experiments in which 10 µM 3β5αP is added to membranes that have been fully equilibrated with a low concentration (3 nM) of [^3^H]muscimol and binding is measured as a function of time. Enhancement of [^3^H]muscimol binding by 10 µM 3β5αP is slow, with a time constant of 4 min at 4 °C (τ = 3.97 ± 0.15 min: mean ± SEM, *n* = 4, Supplementary file 1). In contrast, when α_1_β_3_ GABA_A_Rs are exposed to long pulses of a high concentration of GABA, KK148- and 3β5αP-induced desensitization is rapid (Figures 2A and 2C), since in these conditions almost all of the receptors are either in an open or desensitized conformation and desensitization is not slowed by the transition from resting to open state (57). Thus, the slow enhancement of [^3^H]muscimol binding by 3β5αP (Supplementary file 1) is likely rate-limited by the transition of receptors to activated then to desensitized states at 3 nM muscimol rather than by 3β5αP binding. These time course experiments are most consistent with a model in which receptors preferentially progress from the resting to active to desensitized states, which are then stabilized by the NAM-NS.

The selective binding of 3β5αP to a subset of identified NS binding sites provides an explanation for its NAM activity. 3β5αP stabilizes desensitized receptors by binding to the α_1_ and β_3_ intrasubunit sites, but does not activate the receptor because it does not bind to the intersubunit site. This site-selective binding is unexpected for several reasons. First, docking and free energy perturbation calculations in a prior study predicted that epi-pregnanolone (3β5βP) binds to the intersubunit site in a similar orientation and with free energies of binding that are equivalent to pregnanolone (3α5βP) (23). The modeling suggested that 3β5βP does not form a hydrogen bond with αQ242, a possible explanation for its lack of efficacy (23). Our docking studies also show similar binding energies and orientations of 3β5αP and 3α5αP binding in the β_3_(+)–α_1_(-) intersubunit site (Supplementary file 2). We have also shown that binding affinity or docking scores of NS binding to the intersubunit site is not significantly affected by mutations (α_1_Q242L, α_1_Q242W, α_1_W246L) that eliminate NS activation, although binding orientation is altered (28). These data indicate that NS binding in the intersubunit site is tolerant to significant changes in critical residues and NS ligand structure, and are consistent with our findings that NS analogues, such as KK148 and KK150, can bind to the intersubunit site but have no effect on activation (Figures 1 and 4). Thus, the peculiar lack of 3β5αP binding to the intersubunit site suggests that either: (1) details in the structure of the intersubunit site in the open conformation that explain the absence of 3β5αP binding are not apparent in current high-resolution structures or; (2) 3β5αP does not bind for other reasons. One plausible explanation is that 3β5αP, like cholesterol, has low chemical activity in the membrane and does not achieve sufficiently available concentration to bind in this site (58). This explanation would require that the chemical activity of 3β5αP differs between the inner and outer leaflets of a plasma membrane since 3β5αP binds to the intrasubunit sites.

The functional analysis of mutations in each of the three NS binding sites demonstrates that the activating and desensitizing effects of NS result from occupancy of distinct sites. In particular, binding of certain NS (3β5αP, KK148) to α_1_ and β_3_ intrasubunit sites modulates the open-desensitized equilibrium. Interestingly, lipid binding to intrasubunit pockets in bacterial pLGICs analogous to the α_1_ and β_3_ intrasubunit sites in GABA_A_R, also modulates receptor desensitization; docosahexaenoic acid binding to an intrasubunit site in GLIC (59) and phosphatidylglycerol in ELIC (60) increase and decrease agonist-induced desensitization, respectively. The combined results of mutational analyses and binding data demonstrate that the effects of various NS analogues are also a consequence of their efficacy at each of the sites they occupy. For example, KK148 and KK150 occupy the intersubunit site (Figure 4C), but do not activate GABA_A_R currents (Figures 1C-D), and KK150 occupies both intrasubunit sites (Figure 4B and Figure 4–figure supplement 1) but does not desensitize the receptor (Figure 2B).

To explain the effects of the 3-substituted NS analogues, we propose a model in which NS-selective binding at three distinct binding sites on the GABA_A_R preferentially stabilizes specific states (resting, open, desensitized) of the receptor (Figure 10). Orthosteric agonist (GABA or muscimol) binding shifts the equilibrium towards high-affinity states (open and desensitized). 3α5αP allosterically stabilizes the open state through binding to the β_3_–α_1_ intersubunit and α_1_ intrasubunit sites and stabilizes the desensitized state through the β_3_ intrasubunit site (Figure 10A). In contrast, 3β5αP preferentially stabilizes the desensitized state through binding to both intrasubunit sites (Figure 10B). KK148, like 3β5αP, stabilizes the desensitized state by binding to the intrasubunit sites (Figure 10C). KK148 also binds to the intersubunit site, presumably with no state-dependence, since it is neither an agonist nor an inverse-agonist (Figures 1C and 10C). KK150, which neither activates nor desensitizes GABA_A_Rs and is not an inverse agonist, binds to all three sites, again presumably with no-state dependence (Figures 1D and 10D). This model predicts that KK148 should act as a competitive antagonist to PAM-NS at the intersubunit site. Indeed, KK148 reduces 3α5αP potentiation of GABA-elicited currents by about 24% (Supplementary file 3). The model also predicts that KK150 should be a competitive NS antagonist at all three binding sites. Consistent with this prediction, KK150 weakly antagonizes 3α5αP potentiation of GABA-elicited currents (12%, Supplementary file 3) and antagonizes 3α5αP and KK148 enhancement of [^3^H]muscimol binding (Figure 7). The modest antagonism of 3α5αP potentiation observed with KK148 and KK150 is likely the result of these ligands having lower binding affinity than the 3-OH NS.

**FIGURE 10:**
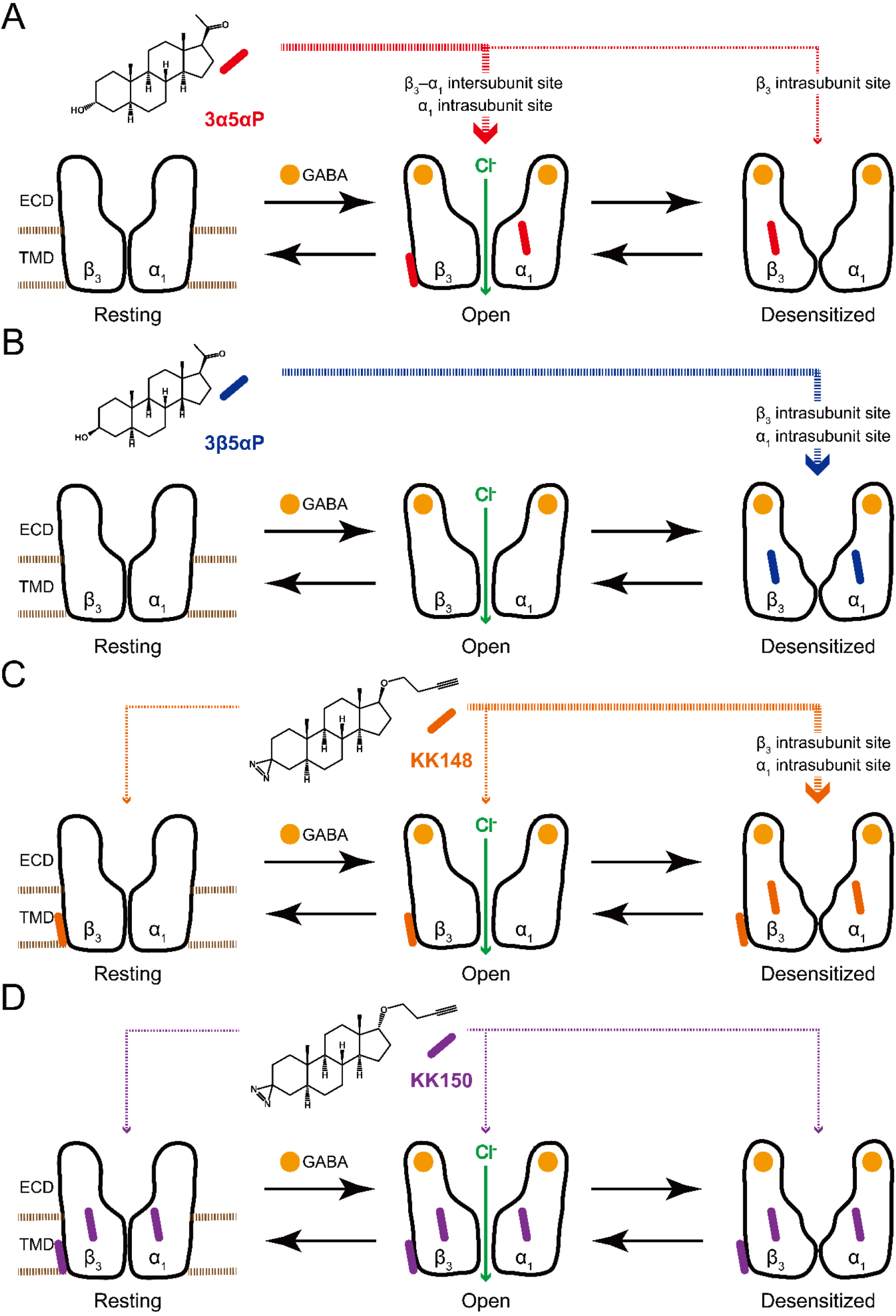
Neurosteroids preferentially stabilize GABA_A_R in different states. (A) Model showing three fundamental conformational states that depict the channel function in the GABA_A_R: a resting state; an open state; and a desensitized state. Agonist (GABA: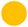) binding shifts the equilibrium towards high-affinity states (open and desensitized). Allopregnanolone (3α5αP: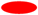) allosterically stabilizes the high-affinity states (an open state through the β_3_–α_1_ intersubunit and the α_1_ intrasubunit sites; a desensitized state through the β_3_ intrasubunit site). The width of red arrows indicates relative affinities of 3α5αP for the open or desensitized state of the receptor. (B) Same as (A) for epi-allopregnanolone (3β5αP: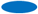). 3β5αP stabilizes a desensitized state through the β_3_ and α_1_ intrasubunit sites. (C) Same as (A) for KK148 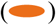. KK148 allosterically stabilizes a desensitized state through the β_3_ and α_1_ intrasubunit sites, and equally stabilizes all three states of the receptor through the β_3_–α_1_ intersubunit site. The width of orange arrows indicates relative affinities of KK148 for each state of the receptor. (D) Same as (A) for KK150 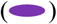. KK150 equally stabilizes all three states of the receptor through the β_3_ and α_1_ intrasubunit sites, and the β_3_–α_1_ intersubunit site.

The site-specific model of NS action (Figure 10) has significant implications for the synaptic mechanisms of PAM-NS action. At a synapse, GABA_A_Rs are transiently exposed to high (mM) concentrations of GABA leading to a channel P_open_ approaching one (61,62). GABA is quickly cleared from the synapse leading to rapid deactivation with minimal desensitization (57,63). In the presence of a PAM-NS, deactivation is slowed, resulting in a prolongation of the inhibitory postsynaptic current (IPSC) and increased inhibitory current (18,55,64,65). This effect is largely attributable to stabilization of the open state, presumably by binding to the intersubunit and α_1_ intrasubunit binding sites. A second effect has been observed in which the PAM-NS 3α5αP (54) and 3α5α-THDOC (55) prolong the slow component of GABA_A_R desensitization and slow recovery from desensitization. This results in increased late channel openings (55,57) and IPSC prolongation (64,65). When the frequency of synaptic firing is rapid, the desensitizing effect of NS may also contribute to frequency-dependent reduction in IPSC amplitude (55,66). The desensitizing effect of 3α5αP is predominantly mediated by binding at the β_3_ intrasubunit site. The balance between stabilization of the open and desensitized channels should be determined by the relative occupancies for the intersubunit site of the active receptor and the β_3_ intrasubunit site of the desensitized receptor. Computational docking of 3α5αP to these sites indicates modest differences in affinity between the sites with a rank order affinity of: intersubunit > α_1_ intrasubunit > β_3_ intrasubunit sites (Supplementary file 2) (11). Mutational analysis of the effects of NS on enhancement of [^3^H]muscimol binding also indicates that 3α5αP has a lower affinity to the β_3_ intrasubunit site (Figure 6B, Table 1). Thus binding to the β_3_ intrasubunit site may serve as a negative feedback mechanism preventing excessive PAM-NS effects on synaptic currents.

In summary, this study describes a unique NS pharmacology in which different NS analogues selectively bind to subsets of three sites on the α_1_β_3_ GABA_A_R, with each analogue exhibiting state-dependent binding at a given site. The combination of site-selectivity and state-dependence of binding determines whether a NS analogue is a PAM, a NAM or an antagonist of NS action at the GABA_A_R. It seems likely that other GABA_A_R subunit isoforms and heteropentameric subunit combinations will reveal additional NS binding sites with distinct affinity and state-dependence for various analogues. The identification of potent agonists and antagonists for each of these sites will provide tools for understanding the biological effects of endogenous neurosteroids and potentially for the development of precision neurosteroid therapeutics.

## ACKNOWLEDGEMENTS

The authors thank Drs. Charles F. Zorumski, Steven Mennerick and Joseph Henry Steinbach for insightful advice and valuable suggestions.

## MATERIALS AND METHODS

### Construct design

The human α_1_ and β_3_ subunits were subcloned into pcDNA3 for molecular manipulations and cRNA synthesis. Using QuikChange mutagenesis (Agilent Technologies, Santa Clara, CA), a FLAG tag was first added to the α_1_ subunit then an octa-histidine tag was added to generate the following His-FLAG tag tandem (QPSLHHHHHHHHDYKDDDDKDEL), inserted between the 4th and 5th residues of the mature peptide. The α_1_ and β_3_ subunits were then transferred into the pcDNA4/TO and pcDNA5/TO vectors (Thermo Fisher Scientific), respectively, for tetracycline inducible expression. Point mutations were generated using the QuikChange site-directed mutagenesis kit and the coding region fully sequenced prior to use. The cDNAs were linearized with Xba I (NEB Labs, Ipswich, MA), and the cRNAs were generated using T7 mMessage mMachine (Ambion, Austin, TX).

### Cell culture and generation of stable cell line

Cell culture was performed as described in previous reports (11). The tetracycline inducible cell line T-REx™-HEK293 (Thermo Fisher Scientific) was cultured under the following conditions: cells were maintained in DMEM/F-12 50/50 medium containing 10% fetal bovine serum (tetracycline-free, Takara, Mountain View, CA), penicillin (100 units/ml), streptomycin (100 g/ml), and blasticidin (2 μg/ml) at 37 °C in a humidified atmosphere containing 5% CO_2_. Cells were passaged twice each week, maintaining subconfluent cultures. Stably transfected cells were cultured as above with the addition of hygromycin (50 μg/ml) and zeocin (20 μg/ml). A stable cell line was generated by transfecting T-REx™-HEK293 cells with human α_1_-8xHis-FLAG pcDNA4/TO and human β_3_ pcDNA5/TO, in a 150 mm culture dish, using the Effectene transfection reagent (Qiagen). Two days after transfection, selection of stably transfected cells was performed with hygromycin and zeocin until distinct colonies appeared. Medium was exchanged several times each week to maintain antibiotic selection. Individual clones were selected from the dish and transferred to 24-well plates for expansion of each clone selected. When the cells grew sufficiently, about 50% confluency, they were split into two other plates, one for a surface ELISA against the FLAG epitope and a second for protein assay, to normalize surface expression to cell number. The best eight clones were selected for expansion into 150 mm dishes, followed by [^3^H]muscimol binding to examine the receptor density. Once the best expressing clone was determined, the highest expressing cells of that clone were selected through fluorescence activated cell sorting.

### Membrane protein preparation

Stably transfected cells were plated into dishes. After reaching 50% confluency, GABA receptors were expressed by inducing cells with 1 μg/ml of doxycycline with the addition of 5 mM sodium butyrate. Cells were harvested 48 to 72 hours after induction. HEK cells, after induction, grown to 100% confluency were harvested and washed with 10 mM potassium phosphate, 100 mM potassium chloride (pH 7.5) plus protease inhibitors (Sigma) two times. The cells were collected by centrifugation at 1,000 g at 4 °C for 5 min. The cells were homogenized with a glass mortar and a Teflon pestle for ten strokes on ice. The pellet containing the membrane proteins was collected after centrifugation at 20,000 g at 4 °C for 45 min and resuspended in a buffer containing 10 mM potassium phosphate, 100 mM potassium chloride (pH 7.5). The protein concentration was determined with micro-BCA protein assay and stored at -80 °C.

### Photolabeling and purification of α_1_β_3_ GABA_A_R

The syntheses of neurosteroid photolabeling reagents (KK148, KK150, KK200, KK123) are detailed in previous reports (33,36). For all the photolabeling experiments, 10-20 mg of HEK cell membrane proteins (about 300 pmol [^3^H]muscimol binding) were thawed and resuspended in buffer containing 10 mM potassium phosphate, 100 mM potassium chloride (pH 7.5) and 1 mM GABA at a final concentration of 1.25 mg/ml. For the photolabeling competition experiments, 3 μM KK200 or KK123 in the presence of 30 μM competitor (3α5αP, KK148, KK150, and 3β5αP) or the same volume of ethanol was added to the membrane proteins and incubated on ice for 1 h. The samples were then irradiated in a quartz cuvette for 5 min, by using a photoreactor emitting light at > 320 nm. The membrane proteins were then collected by centrifugation at 20,000 g at 4 °C for 45 min. The photolabeled membrane proteins were resuspended in lysis buffer containing 1% *n*-dodecyl-β-D-maltoside (DDM), 0.25% cholesteryl hemisuccinate (CHS), 50 mM Tris (pH 7.5), 150 mM NaCl, 2 mM CaCl_2_, 5 mM KCl, 5 mM MgCl_2_, 1 mM EDTA, 10% glycerol at a final concentration of 1 mg/ml. The membrane protein suspension was homogenized using a glass mortar and a Teflon pestle and incubated at 4 °C overnight. The protein lysate was centrifuged at 20,000 g at 4 °C for 45 min and supernatant was incubated with 0.5 ml anti-FLAG agarose (Sigma) at 4 °C for 2 h. The anti-FLAG agarose was then transferred to an empty column, followed by washing with 20 ml washing buffer (50 mM triethylammonium bicarbonate and 0.05% DDM). The GABA_A_Rs were eluted with aliquots of 200 μg/ml FLAG tag peptide and 100 μg/ml 3X FLAG (ApexBio) in the washing buffer. The pooled eluates (9 ml) containing GABA_A_Rs were concentrated to 100 μl using 100 kDa cut-off centrifugal filters.

### Tryptic middle-down MS analysis

The purified α_1_β_3_ GABA_A_R (100 μl) was reduced with 5 mM tris(2-carboxyethyl)phosphine for 1 h, alkylated with 5 mM *N*-ethylmaleimide (NEM) for 1 h, and quenched with 5 mM dithiothreitol (DTT) for 15 min. These three steps were done at RT. Samples were then digested with 8 μg of trypsin for 7 days at 4 °C to obtain maximal recovery of TMD peptides. Next, the digestion was terminated by adding formic acid in a final concentration of 1%, followed directly by LC-MS analysis on an Orbitrap Elite mass spectrometer. 20 μl samples were injected into a home-packed PLRP-S (Agilent) column (10 cm x 75 μm, 300 Å), separated with a 145 min gradient from 10 to 90% acetonitrile, and introduced to the mass spectrometer at 800 nl/min with a nanospray source. MS acquisition was set as a MS1 Orbitrap scan (resolution of 60,000) followed by top 20 MS2 Orbitrap scans (resolution of 15,000) using data-dependent acquisition, and exclusion of singly charged precursors. Fragmentation was performed using high-energy dissociation with normalized energy of 35%. Analysis of data sets was performed using Xcalibur (Thermo Fisher Scientific) to manually search for TM1, TM2, TM3 or TM4 tryptic peptides with or without neurosteroid photolabeling modifications. Photolabeling efficiency was estimated by generating extracted chromatograms of unlabeled and labeled peptides, determining the area under the curve, and calculating the abundance of labeled peptide/(unlabeled + labeled peptide). Analysis of statistical significance comparing the photolabeling efficiency of KK200 and KK123 for α_1_β_3_ GABA_A_R was determined using one-way ANOVA with *post hoc* Bonferroni correction and paired *t*-test, respectively (GraphPad Prism 6). MS2 spectra of photolabeled TMD peptides were analyzed by manual assignment of fragment ions with and without photolabeling modification. Fragment ions were accepted based on the presence of a monoisotopic mass within 20 ppm mass accuracy. In addition to manual analysis, PEAKS (Bioinformatics Solutions Inc.) database searches were performed for data sets of photolabeled α_1_β_3_ GABA_A_R. Search parameters were set for a precursor mass accuracy of 20 ppm, fragment ion accuracy of 0.1 Da, up to 3 missed cleavages on either end of the peptide, false discovery rate of 0.1%, and variable modifications of methionine oxidation, cysteine alkylation with NEM and DTT, and NS analogue photolabeling reagents on any amino acid.

### Radioligand binding assays

[^3^H]muscimol binding assays were performed using a previously described method (11). HEK cell membrane proteins (100 μg/ml final concentration) were incubated with 3 nM [^3^H]muscimol (30 Ci/mmol; PerkinElmer Life Sciences), neurosteroid (3 nM–30 μM) or etomidate (30 nM–200 μM) in different concentrations, binding buffer (10 mM potassium phosphate, 100 mM potassium chloride, pH 7.5), in a total volume of 1 ml. Assay tubes were incubated for 1 h on ice in the dark. Nonspecific binding was determined by binding in the presence of 1 mM GABA. Membranes were collected on Whatman/GF-C glass filter papers using a Brandel cell harvester (Gaithersburg, MD). To perform [^3^H]muscimol binding isotherms, 100 μg/ml aliquots of membrane protein were incubated with 0.3 nM–1 μM [^3^H]muscimol, with specific activity reduced to 2 Ci/mmol, for 1 h on ice in the dark. The membranes were collected on Whatman/GF-B glass filter paper using a vacuum manifold. For the [^3^H]muscimol binding competition, neurosteroids in the presence of competitors (0.3 μM 3α5αP or 3 μM KK148 vs. 30 μM KK150) or the same volume of dimethyl sulfoxide (DMSO) were added to the membrane proteins and incubated with 3 nM [^3^H]muscimol on ice for 1 h. Time courses of neurosteroid [^3^H]muscimol binding enhancement were examined by adding 10 µM of neurosteroids (3α5αP, 3β5αP) to membranes that had been fully equilibrated with 3 nM [^3^H]muscimol for 1 h on ice and binding was measured as a function of time at 1, 3, 10, 30, 60 min. The membranes were collected on Whatman/GF-B glass filter papers using vacuum manifold. Radioactivity bound to the filters was measured by liquid scintillation spectrometry using Bio-Safe II (Research Products International Corporation).

### Radioligand binding to intact cells

HEK cells were harvested by gently washing dishes with buffer containing 10 mM sodium phosphate (pH 7.5), 150 mM sodium chloride twice. The cells were collected by centrifugation at 500 g at 4 °C for 5 min, and resuspended in isotonic (10 mM sodium phosphate, 150 mM sodium chloride, pH 7.5) or hypotonic (10 mM sodium phosphate, pH 7.5) buffer to prepare two different conditions for radioligand binding to intact cells [isotonic buffer for cell surface receptors; hypotonic buffer for total receptors (cell surface receptors + intracellular receptors)]. The cells were incubated on ice for 2 h, after which the sodium chloride concentration was adjusted to be isotonic before the radioligand binding procedure. HEK cells were aliquoted to assay tubes (20 samples/150 mm dish) in a total volume of 1 ml, and incubated with 3 nM [^3^H]muscimol and 10 μM KK148 for 1 h on ice in the dark. Nonspecific binding was determined by binding in the presence of 1 mM GABA. The membranes were collected on Whatman/GF-B glass filter papers using vacuum manifold. Radioactivity bound to the filters was measured by liquid scintillation spectrometry using Bio-Safe II.

### Receptor expression in Xenopus laevis oocytes and electrophysiological recordings

The wild-type and mutant α_1_β_3_ GABA_A_R were expressed in oocytes from the African clawed frog (*Xenopus laevis*). Frogs were purchased from Xenopus 1 (Dexter, MI), and housed and cared for in a Washington University Animal Care Facility under the supervision of the Washington University Division of Comparative Medicine. Harvesting of oocytes was conducted under the Guide for the Care and Use of Laboratory Animals as adopted and promulgated by the National Institutes of Health. The animal protocol (No. 20180191) was approved by the Animal Studies Committee of Washington University in St. Louis.

The oocytes were injected with a total of 12 ng cRNA. The ratio of cRNAs was 5:1 ratio (α_1_:β_3_) to minimize the expression of β_3_ homomeric receptors. Following injection, the oocytes were incubated in ND96 (96 mM NaCl, 2 mM KCl, 1.8 mM CaCl_2_, 1 mM MgCl_2_, 5 mM HEPES; pH 7.4) with supplements (2.5 mM Na pyruvate, 100 U/ml penicillin, 100 μg/ml streptomycin and 50 μg/ml gentamycin) at 16 °C for 2 days prior to conducting electrophysiological recordings. The electrophysiological recordings were conducted at room temperature using standard two-electrode voltage clamp. The oocytes were clamped at -60 mV. The chamber (RC-1Z, Warner Instruments, Hamden, CT) was perfused with ND96 at 5-8 ml/min. Solutions were gravity-applied from 30 ml glass syringes with glass luer slips via Teflon tubing. The current responses were amplified with an OC-725C amplifier (Warner Instruments, Hamden, CT), digitized with a Digidata 1200 series digitizer (Molecular Devices), and stored using pClamp (Molecular Devices). Current traces were analyzed with Clampfit (Molecular Devices). The stock solution of GABA was made in ND96 at 500 mM, stored in aliquots at -20 °C, and diluted on the day of experiment. The stock solution of muscimol was made at 20 mM in ND96 and stored at 4 °C. The steroids were dissolved in DMSO at 10-20 mM and stored at room temperature.

### Electrophysiological data analysis and simulations

The raw amplitudes of the current traces were converted to units of open probability through comparison to the peak response to 1 mM GABA + 50 µM propofol, that was considered to have a peak P_open_ indistinguishable from 1 (11). The level of constitutive activity in the absence of any applied agonist was considered negligible and not included in this calculation. The converted current traces were analyzed in the framework of the three-state Resting-Open-Desensitized activation model (44,45). The model enables analysis and prediction of peak responses using four parameters that characterize the extent of constitutive activity (termed L; L = Resting/Open), affinity of the resting receptor to agonist (K_C_), affinity of the open receptor to agonist (K_O_), and the number of agonist binding sites (N). Analysis and prediction of steady-state responses requires an additional parameter, termed Q, that describes the equilibrium between open and desensitized receptors (Q = Open/Desensitized).

The P_open_ of the peak response is expressed as:

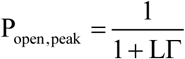

and the P_open_ of the steady-state response as:

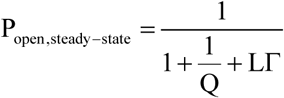

where

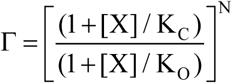

[X] is the concentration of agonist present, and other terms are as described above. In practice, the value of LΓ was calculated using the experimentally-determined P_open_ of the peak response, and then used as a fixed value in estimating Q from P_open,steady-state_.

The P_desensitized_ was calculated using:

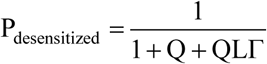

The effect of 3β5αP on 1 mM GABA-elicited steady-state current was expressed through a change in the value of Q. The modified Q (termed Q*) was then used to predict changes in P_open_ and P_desensitized_ at low [GABA].

### Docking simulations

A docking template of the α_1_β_3_ GABA_A_R was developed using the crystal structure of the human β_3_ homopentamer (PDB ID: 4COF) (67). In this structure, the cytoplasmic loop was replaced with the sequence SQPARAA (68). The pentamer subunits were organized as A α_1_, B β_3_, C α_1_, D β_3_, E β_3_. The α_1_ sequence was aligned to the β_3_ sequence using the program MUSCLE (69). The pentameric alignment was then used as input for the program Modeller (70), using 4COF as the template; a total of 25 models were generated. The best model as evaluated by the DOPE score (71) was then submitted to the H++ server (http://biophysics.cs.vt.edu) to determine charges and optimize hydrogen bonding. The optimized structure was then submitted to the PPM server (https://opm.phar.umich.edu/ppm_server) for orientation into a lipid membrane. The correctly oriented receptor was then submitted to the CHARMM-GUI Membrane Builder server (http://www.charmm-gui.org) to build the fully solvated, membrane bound system oriented into a 1-palmitoyl-2-oleoyl-sn-glycero-3-phosphatidylcholine (POPC) bilayer. The system was fully solvated with 40715 TIP3 water molecules and ionic strength set to 0.15 M KCl. The NAMD input files produced by CHARMM-GUI (72) use a seven-step process of gradually loosening constraints in the simulation prior to production runs. A 100 ns molecular dynamics trajectory was then obtained using the CHARMM36 force field and NAMD (72). The resulting trajectory was then processed using the utility mdtraj (73), to extract a snapshot of the receptor at each nanosecond of time frame. These structures were then mutually aligned by fitting the alpha carbons, providing a set of 100 mutually aligned structures used for docking studies. The docking was performed using AutoDock Vina (74) on each of the 100 snapshots in order to capture receptor flexibility. 3α5αP and 3β5αP were prepared by converting the sdf file from PubChem into a PDB file using Open Babel (75), and Gasteiger charges and free torsion angles were determined by AutoDock Tools. Docking grid boxes were built for the β_3_-α_1_ intersubunit, and the α_1_ and β_3_ intrasubunit sites with dimensions of 15 × 15 × 15 Ångströms encompassing each binding pocket. Docking was limited to an energy range of 3 kcal from the best docking pose and was limited to a total of 20 unique poses. The docking results for a given site could result in a maximum of 2,000 unique poses (20 poses × 100 receptor structures); these were then clustered geometrically using the program DIVCF (76). The resulting clusters were ranked by Vina score and cluster size, and then visually analyzed.

## SUPPLEMENTAL MATERIALS

**FIGURE 1–FIGURE SUPPLEMENT 1:**
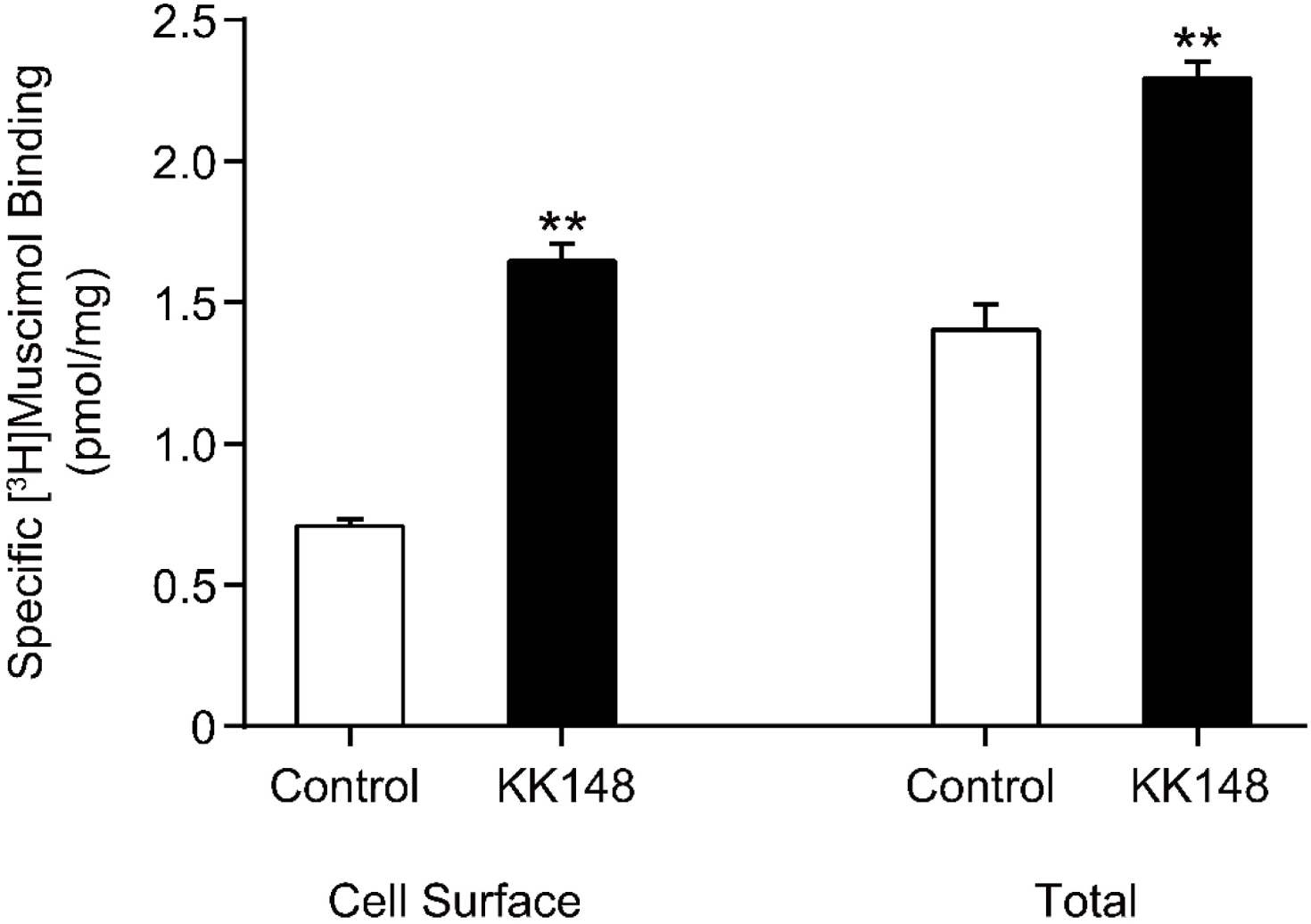
Neurosteroid modulation of muscimol binding to intact cells. Enhancement of specific [^3^H]muscimol (3 nM) binding to α_1_β_3_ GABA_A_Rs on intact HEK cell surfaces (left bars) and total receptors (cell surface receptors + intracellular receptors, right bars) by 10 μM KK148 (*n* = 6, ± SEM). The larger amount of control binding in total vs. cell surface demonstrates the distribution of GABA_A_R between plasma membrane and intracellular membrane. \*\**P* < 0.01 vs. control.

**FIGURE 4–FIGURE SUPPLEMENT 1:**
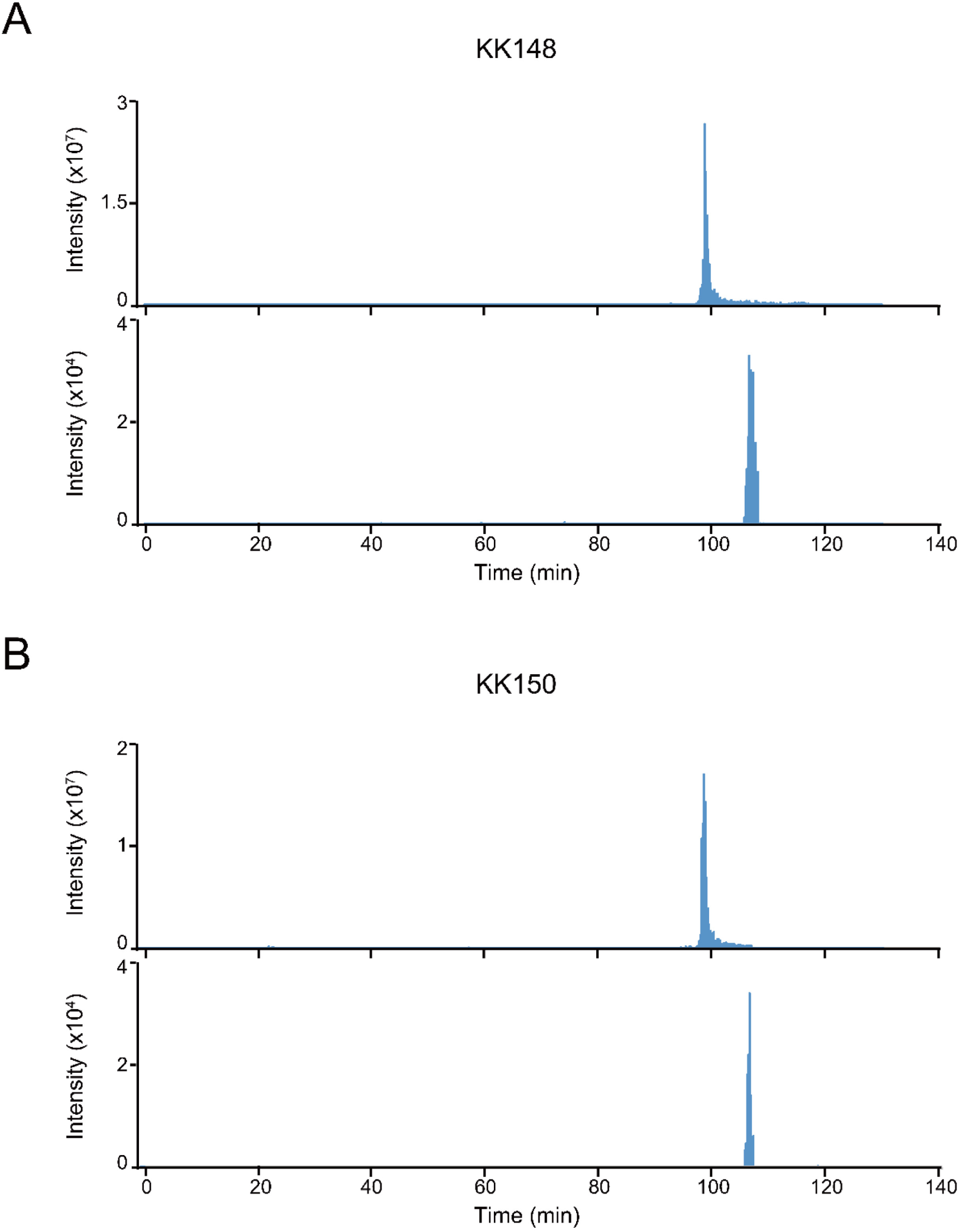
Extracted ion chromatograms of labeled and unlabeled β_3_ subunit TM4 peptides. (A) Extracted ion chromatogram (XIC) of the β_3_ subunit TM4 tryptic peptide in the α_1_β_3_ GABA_A_R. The upper and lower XIC show representative unlabeled β_3_ subunit TM4 peptide and the peptide labeled with 30 μM KK148, respectively. (B) Same as (A) for KK150.

**FIGURE 4–FIGURE SUPPLEMENT 2:**
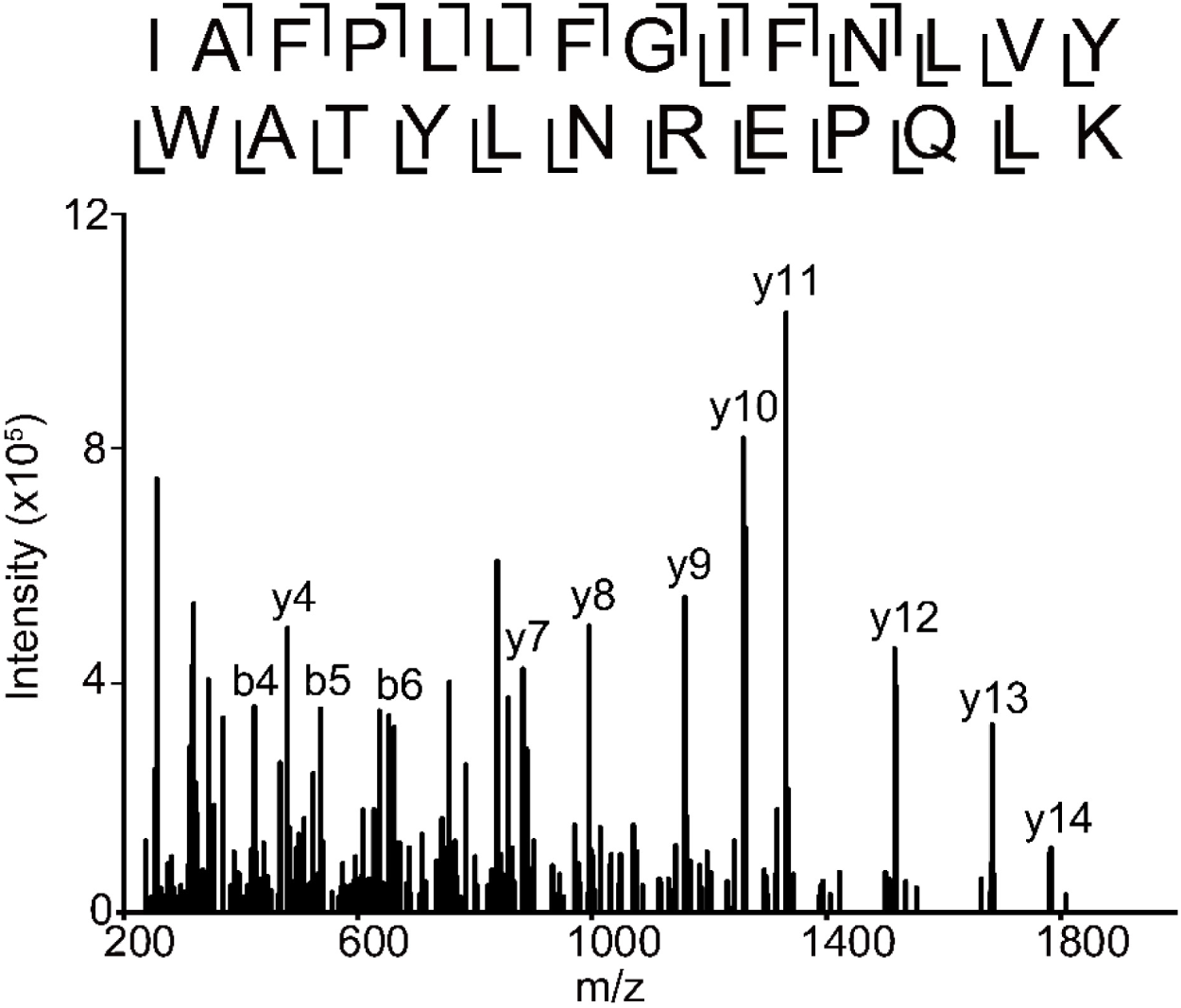
Fragmentation spectrum of unlabeled α_1_ subunit TM4 peptide. HCD fragmentation spectrum of the α_1_ subunit TM4 unlabeled tryptic peptide in the α_1_β_3_ GABA_A_R.

**FIGURE 6–FIGURE SUPPLEMENT 1:**
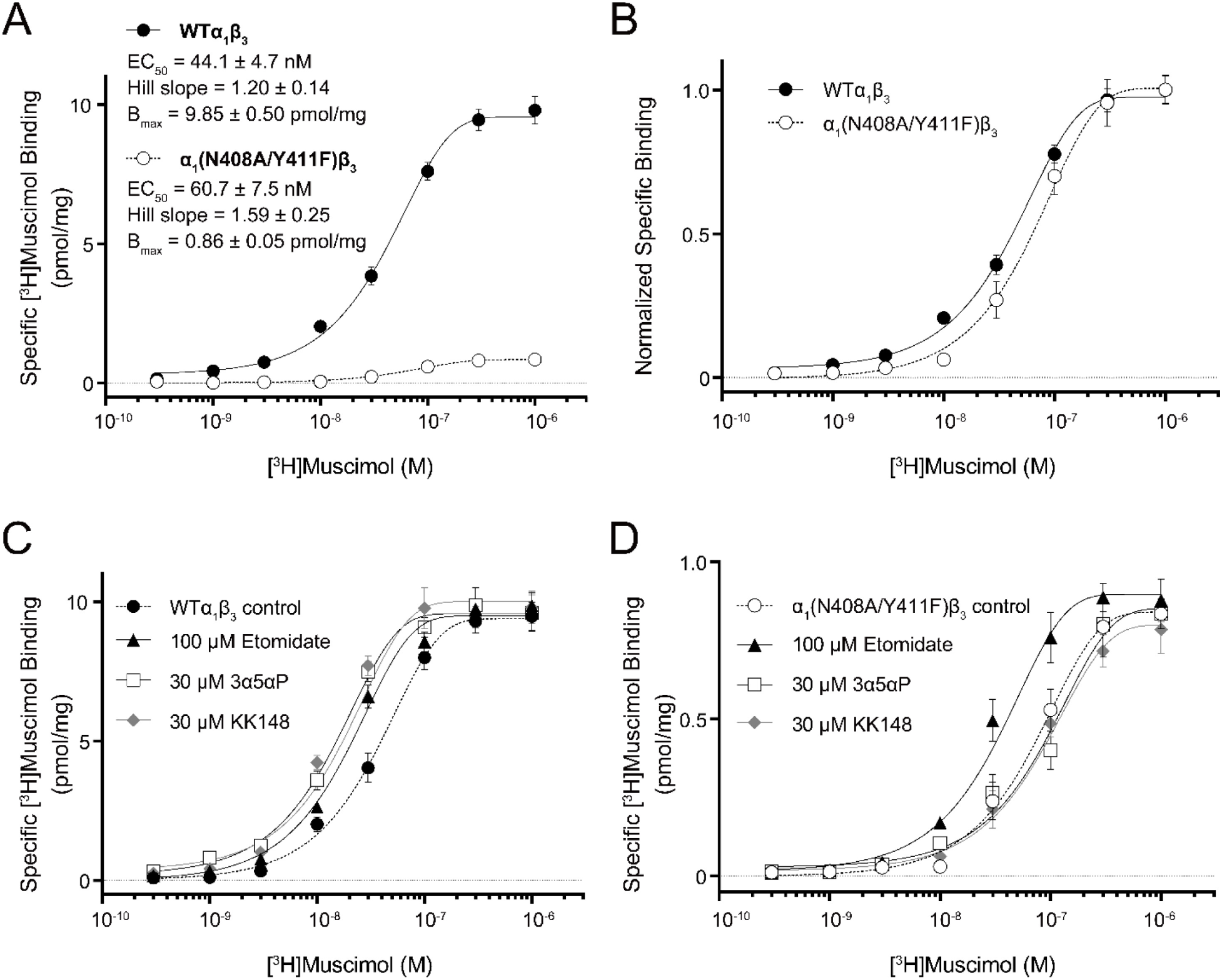
Neurosteroid effect on [^3^H]muscimol binding isotherms in α_1_β_3_ WT and α_1_(N408A/Y411F)β_3_ GABA_A_Rs. (A) [^3^H]muscimol binding isotherms (0.3 nM–1 μM) for α_1_β_3_ GABA_A_R WT and α_1_(N408A/Y411F)β_3_ GABA_A_R. The EC_50_, Hill slope and B_max_ are presented as mean ± SEM (*n* = 3). (B) Normalized curves of specific [^3^H]muscimol binding shown in (A). (C) Effect of 100 μM etomidate, 30 μM allopregnanolone (3α5αP) and 30 μM KK148 on [^3^H]muscimol binding isotherms in the α_1_β_3_ GABA_A_R WT. (D) Same as (C) in the α_1_(N408A/Y411F)β_3_ mutant. Each data point represents mean ± SEM from triplicate experiments.

**FIGURE 6–FIGURE SUPPLEMENT 2:**
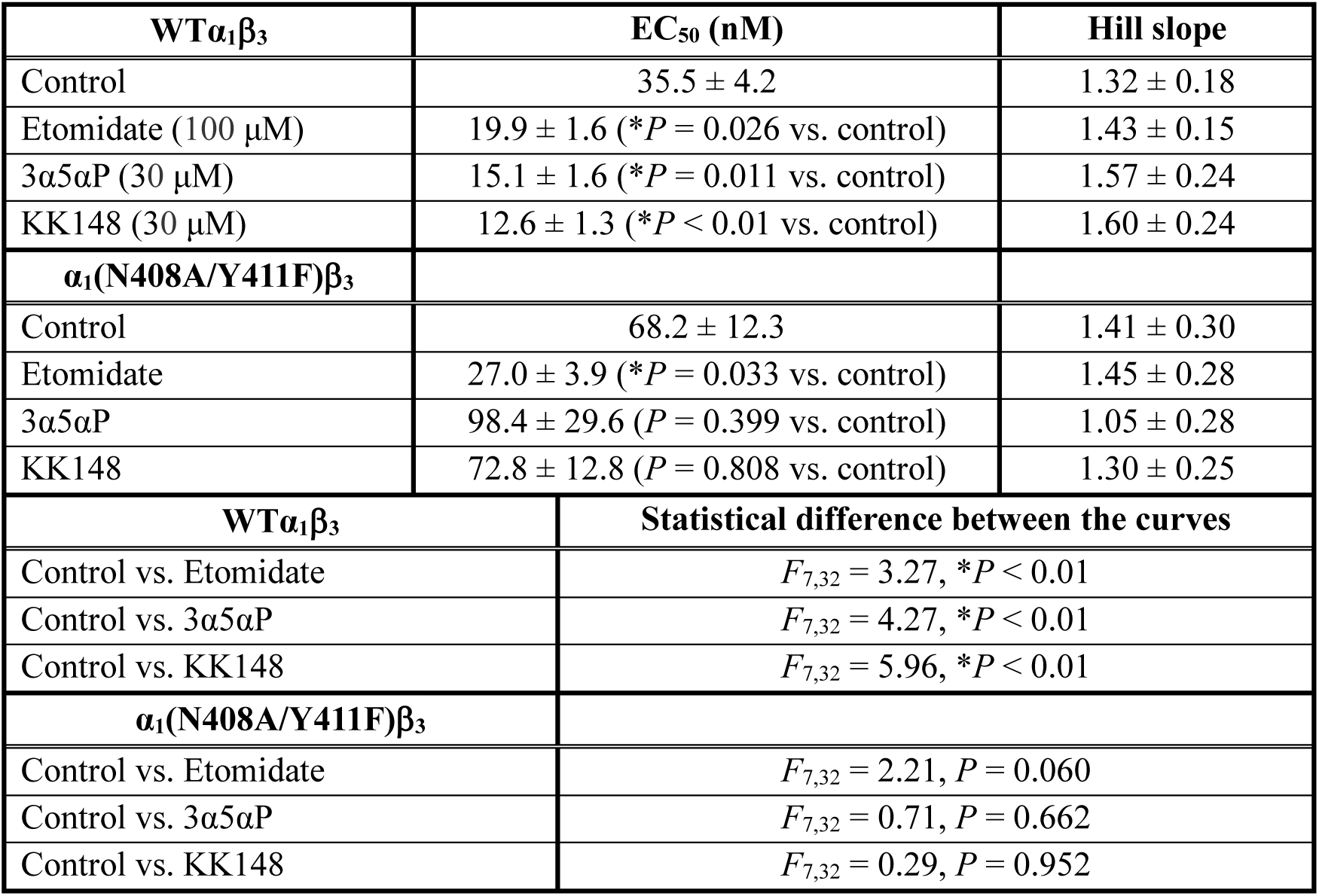
Properties of [^3^H]muscimol saturation binding curves in the α_1_β_3_ GABA_A_R WT and α_1_(N408A/Y411F)β_3_ mutant. EC_50_ and Hill slope for the [^3^H]muscimol saturation binding curves in Figure 6–figure supplement 1C-D. EC_50_ values were compared by unpaired *t*-test. Statistical differences between the whole curves are analyzed using two-way ANOVA. Data are presented as mean ± SEM (*n* = 3).

**SUPPLEMENTARY FILE 1:**
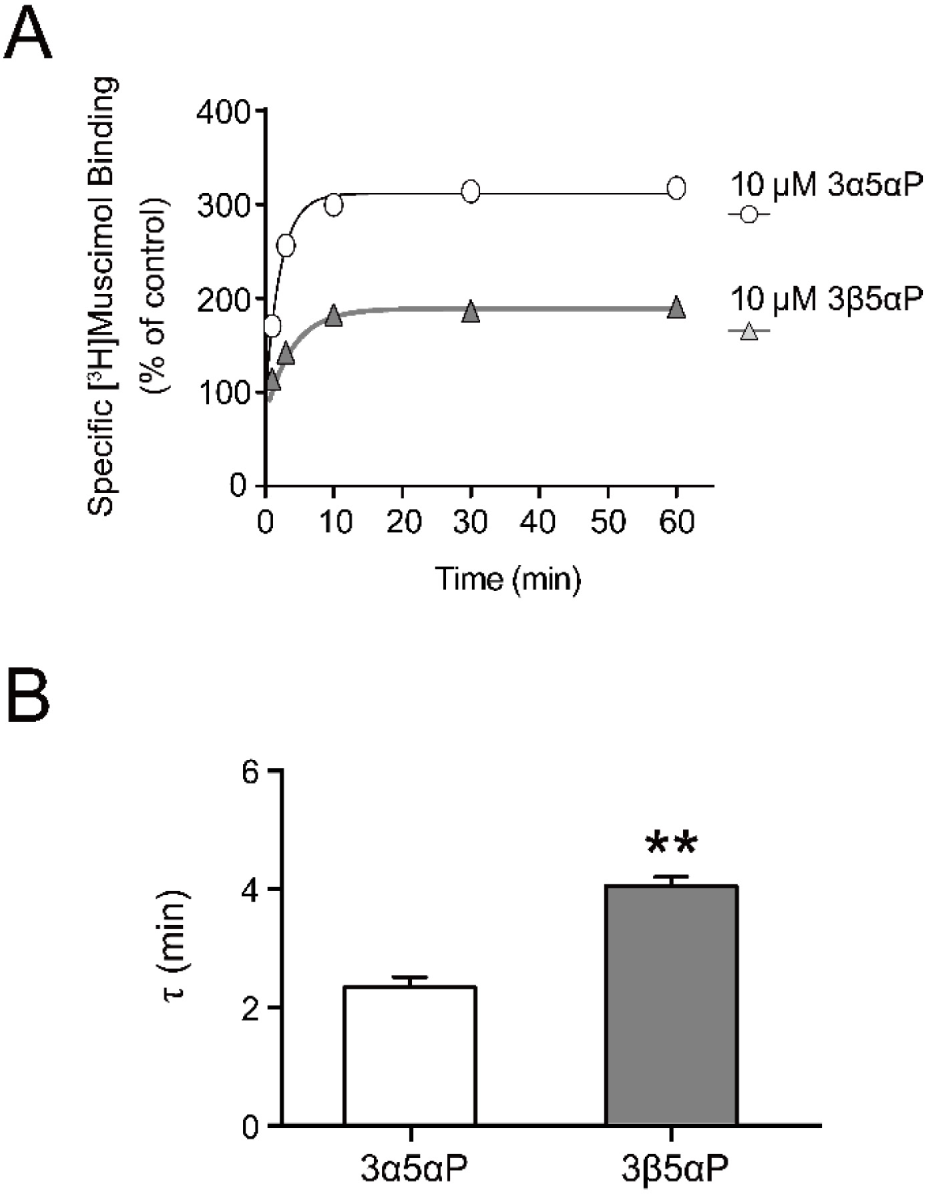
Time course of neurosteroid modulation of muscimol binding. (A) Time course of [^3^H]muscimol binding enhancement by 10 µM allopregnanolone (3α5αP) and epi-allopregnanolone (3β5αP). Neurosteroids were added to α_1_β_3_ GABA_A_R membranes that had been fully equilibrated with 3 nM [^3^H]muscimol and binding was measured as a function of time (*n* = 4, ± SEM). (B) Time constants for neurosteroid-induced enhancement of [^3^H]muscimol binding (*n* = 4, ± SEM). \*\**P* < 0.01 vs. 3α5αP.

**SUPPLEMENTARY FILE 2:**
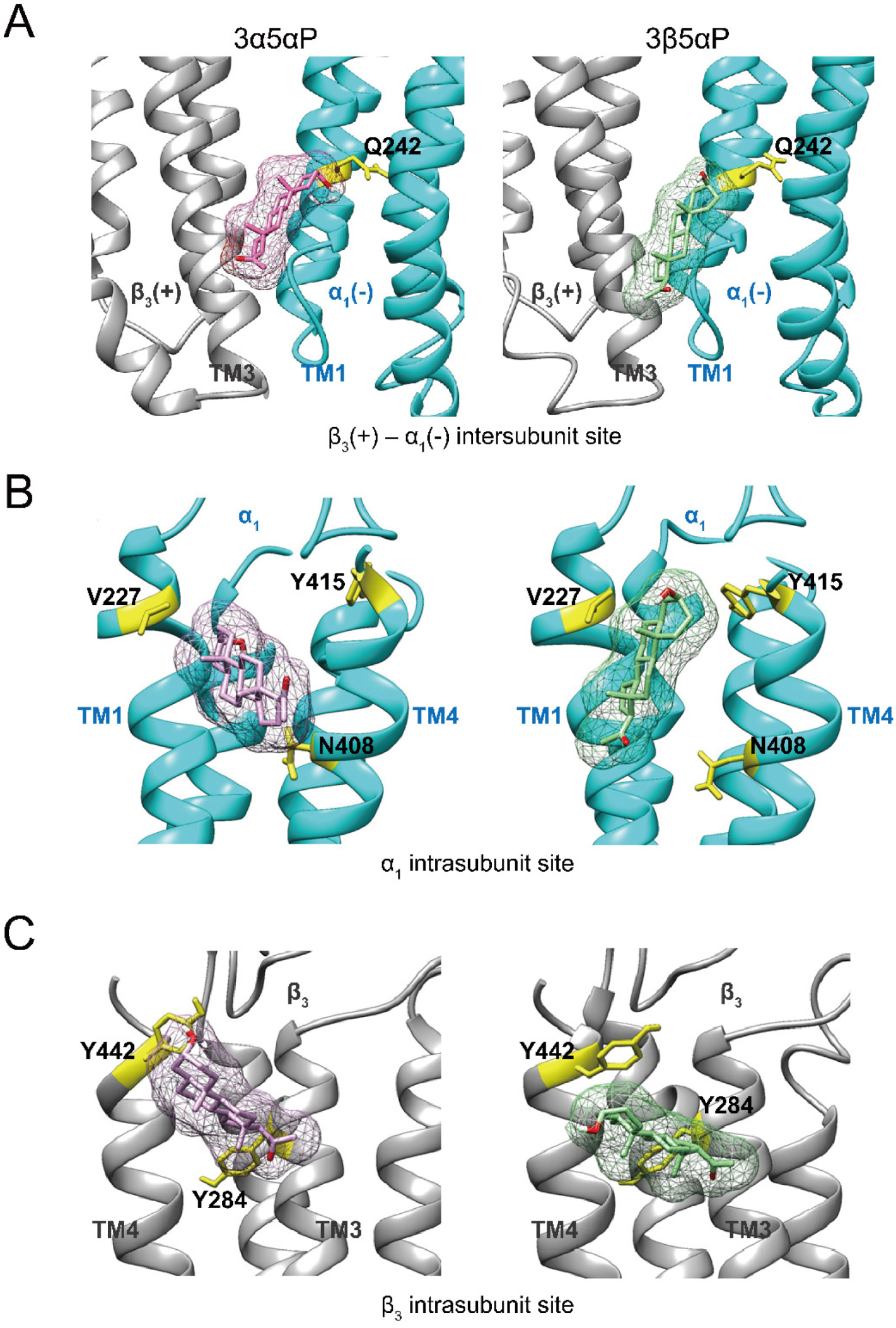
Allopregnanolone and epi-allopregnanolone docking poses within three neurosteroid binding pockets in the α_1_β_3_ GABA_A_R TMD. (A) Representative poses for allopregnanolone (3α5αP) (pink) and epi-allopregnanolone (3β5αP) (light green) docked within the β_3_(+)–α_1_(-) intersubunit site (Vina score: - 7.2 for 3α5αP; -5.9 for 3β5αP). The receptor snapshot is shown along with the α_1_Q242 side chain (yellow). (B) Same as (A) for the α_1_ intrasubunit site (Vina score: -5.6 for 3α5αP; -5.8 for 3β5αP). The receptor snapshot is shown along with the α_1_V227, α_1_Y415 and α_1_N408 side chains. (C) Same as (A) for the β_3_ intrasubunit site (Vina score: -4.0 for 3α5αP; -3.6 for 3β5αP). The receptor snapshot is shown along with the β_3_Y284 and β_3_Y442 side chains.

**SUPPLEMENTARY FILE 3:**
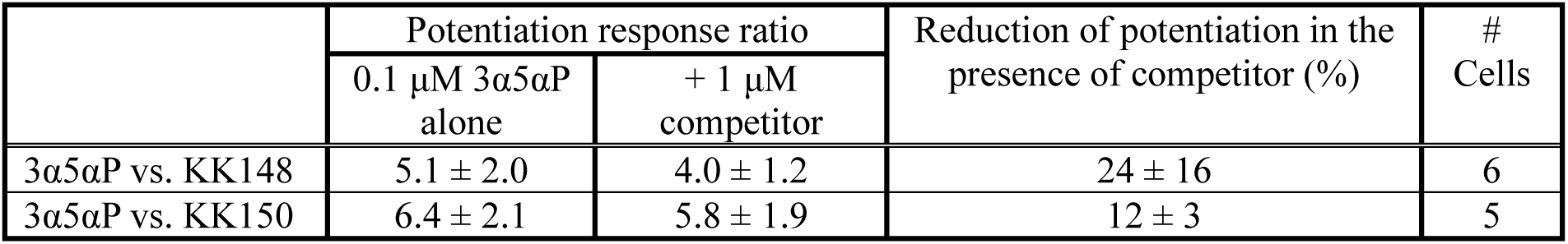
KK148 and KK150 decrease allopregnanolone-induced potentiation. Neurosteroid potentiation of α_1_β_3_ GABA_A_Rs in *Xenopus laevis* oocytes is expressed as potentiation response ratio, calculated as the ratio of the peak responses in the presence of GABA and 0.1 μM allopregnanolone (3α5αP) to the peak response in the presence of GABA alone. The GABA concentrations were selected to generate a response of 3–9% of the response to saturating GABA (30 μM). Co-application of KK148 or KK150 reduces the potentiating effect of 3α5αP on GABA-elicited currents. Data are shown as mean ± SD.

## REFERENCES

1. Belelli, D., and Lambert, J. J. (2005) Neurosteroids: endogenous regulators of the GABA(A) receptor. Nat Rev Neurosci 6, 565–575

2. Mitchell, E. A., Herd, M. B., Gunn, B. G., Lambert, J. J., and Belelli, D. (2008) Neurosteroid modulation of GABAA receptors: molecular determinants and significance in health and disease. Neurochem Int 52, 588–595

3. Represa, A., and Ben-Ari, Y. (2005) Trophic actions of GABA on neuronal development. Trends Neurosci 28, 278–283

4. Grobin, A. C., Gizerian, S., Lieberman, J. A., and Morrow, A. L. (2006) Perinatal allopregnanolone influences prefrontal cortex structure, connectivity and behavior in adult rats. Neuroscience 138, 809–819

5. Akk, G., Covey, D. F., Evers, A. S., Steinbach, J. H., Zorumski, C. F., and Mennerick, S. (2007) Mechanisms of neurosteroid interactions with GABA(A) receptors. Pharmacol Ther 116, 35–57

6. Reddy, D. S., and Estes, W. A. (2016) Clinical Potential of Neurosteroids for CNS Disorders. Trends Pharmacol Sci 37, 543–561

7. Kharasch, E. D., and Hollmann, M. W. (2015) Steroid Anesthesia Revisited: Again. Anesth Analg 120, 983–984

8. Gunduz-Bruce, H., Silber, C., Kaul, I., Rothschild, A. J., Riesenberg, R., Sankoh, A. J., Li, H., Lasser, R., Zorumski, C. F., Rubinow, D. R., Paul, S. M., Jonas, J., Doherty, J. J., and Kanes, S. J. (2019) Trial of SAGE-217 in Patients with Major Depressive Disorder. N Engl J Med 381, 903–911

9. Zorumski, C. F., Paul, S. M., Covey, D. F., and Mennerick, S. (2019) Neurosteroids as novel antidepressants and anxiolytics: GABA-A receptors and beyond. Neurobiol Stress 11, 100196

10. Akk, G., Covey, D. F., Evers, A. S., Mennerick, S., Zorumski, C. F., and Steinbach, J. H. (2010) Kinetic and structural determinants for GABA-A receptor potentiation by neuroactive steroids. Curr Neuropharmacol 8, 18–25

11. Chen, Z. W., Bracamontes, J. R., Budelier, M. M., Germann, A. L., Shin, D. J., Kathiresan, K., Qian, M. X., Manion, B., Cheng, W. W. L., Reichert, D. E., Akk, G., Covey, D. F., and Evers, A. S. (2019) Multiple functional neurosteroid binding sites on GABAA receptors. PLoS Biol 17, e3000157

12. Olsen, R. W. (2018) GABAA receptor: Positive and negative allosteric modulators. Neuropharmacology 136, 10–22

13. Akk, G., Bracamontes, J., and Steinbach, J. H. (2001) Pregnenolone sulfate block of GABA(A) receptors: mechanism and involvement of a residue in the M2 region of the alpha subunit. J Physiol 532, 673–684

14. Wang, M., He, Y., Eisenman, L. N., Fields, C., Zeng, C. M., Mathews, J., Benz, A., Fu, T., Zorumski, E., Steinbach, J. H., Covey, D. F., Zorumski, C. F., and Mennerick, S. (2002) 3beta - hydroxypregnane steroids are pregnenolone sulfate-like GABA(A) receptor antagonists. J Neurosci 22, 3366–3375

15. Shen, W., Mennerick, S., Covey, D. F., and Zorumski, C. F. (2000) Pregnenolone sulfate modulates inhibitory synaptic transmission by enhancing GABA(A) receptor desensitization. J Neurosci 20, 3571–3579

16. Lundgren, P., Stromberg, J., Backstrom, T., and Wang, M. (2003) Allopregnanolone-stimulated GABA-mediated chloride ion flux is inhibited by 3beta-hydroxy-5alpha-pregnan-20-one (isoallopregnanolone). Brain Res 982, 45–53

17. Seljeset, S., Bright, D. P., Thomas, P., and Smart, T. G. (2018) Probing GABA_A_ receptors with inhibitory neurosteroids. Neuropharmacology 136, 23–36

18. Harrison, N. L., Majewska, M. D., Harrington, J. W., and Barker, J. L. (1987) Structure-activity relationships for steroid interaction with the gamma-aminobutyric acidA receptor complex. J Pharmacol Exp Ther 241, 346–353

19. Sigel, E., and Steinmann, M. E. (2012) Structure, function, and modulation of GABA(A) receptors. J Biol Chem 287, 40224–40231

20. Sieghart, W. (2015) Allosteric modulation of GABAA receptors via multiple drug-binding sites. Adv Pharmacol 72, 53–96

21. Olsen, R. W., and Sieghart, W. (2008) International Union of Pharmacology. LXX. Subtypes of gamma-aminobutyric acid(A) receptors: classification on the basis of subunit composition, pharmacology, and function. Update. Pharmacol Rev 60, 243–260

22. Laverty, D., Desai, R., Uchanski, T., Masiulis, S., Stec, W. J., Malinauskas, T., Zivanov, J., Pardon, E., Steyaert, J., Miller, K. W., and Aricescu, A. R. (2019) Cryo-EM structure of the human alpha1beta3gamma2 GABAA receptor in a lipid bilayer. Nature 565, 516–520

23. Miller, P. S., Scott, S., Masiulis, S., De Colibus, L., Pardon, E., Steyaert, J., and Aricescu, A. R. (2017) Structural basis for GABAA receptor potentiation by neurosteroids. Nat Struct Mol Biol 24, 986–992

24. Laverty, D., Thomas, P., Field, M., Andersen, O. J., Gold, M. G., Biggin, P. C., Gielen, M., and Smart, T. G. (2017) Crystal structures of a GABAA-receptor chimera reveal new endogenous neurosteroid-binding sites. Nat Struct Mol Biol 24, 977–985

25. Hosie, A. M., Wilkins, M. E., da Silva, H. M., and Smart, T. G. (2006) Endogenous neurosteroids regulate GABAA receptors through two discrete transmembrane sites. Nature 444, 486–489

26. Hosie, A. M., Clarke, L., da Silva, H., and Smart, T. G. (2009) Conserved site for neurosteroid modulation of GABA A receptors. Neuropharmacology 56, 149–154

27. Chen, Q., Wells, M. M., Arjunan, P., Tillman, T. S., Cohen, A. E., Xu, Y., and Tang, P. (2018) Structural basis of neurosteroid anesthetic action on GABAA receptors. Nat Commun 9, 3972

28. Sugasawa, Y., Bracamontes, J. R., Krishnan, K., Covey, D. F., Reichert, D. E., Akk, G., Chen, Q., Tang, P., Evers, A. S., and Cheng, W. W. L. (2019) The molecular determinants of neurosteroid binding in the GABA(A) receptor. J Steroid Biochem Mol Biol 192, 105383

29. Akk, G., Li, P., Bracamontes, J., Reichert, D. E., Covey, D. F., and Steinbach, J. H. (2008) Mutations of the GABA-A receptor alpha1 subunit M1 domain reveal unexpected complexity for modulation by neuroactive steroids. Mol Pharmacol 74, 614–627

30. Akk, G., Bracamontes, J. R., Covey, D. F., Evers, A., Dao, T., and Steinbach, J. H. (2004) Neuroactive steroids have multiple actions to potentiate GABAA receptors. J Physiol 558, 59–74

31. Evers, A. S., Chen, Z. W., Manion, B. D., Han, M., Jiang, X., Darbandi-Tonkabon, R., Kable, T., Bracamontes, J., Zorumski, C. F., Mennerick, S., Steinbach, J. H., and Covey, D. F. (2010) A synthetic 18-norsteroid distinguishes between two neuroactive steroid binding sites on GABAA receptors. J Pharmacol Exp Ther 333, 404–413

32. Park-Chung, M., Malayev, A., Purdy, R. H., Gibbs, T. T., and Farb, D. H. (1999) Sulfated and unsulfated steroids modulate gamma-aminobutyric acidA receptor function through distinct sites. Brain Res 830, 72–87

33. Jiang, X., Shu, H. J., Krishnan, K., Qian, M., Taylor, A. A., Covey, D. F., Zorumski, C. F., and Mennerick, S. (2016) A clickable neurosteroid photolabel reveals selective Golgi compartmentalization with preferential impact on proximal inhibition. Neuropharmacology 108, 193–206

34. Budelier, M. M., Cheng, W. W. L., Bergdoll, L., Chen, Z. W., Janetka, J. W., Abramson, J., Krishnan, K., Mydock-McGrane, L., Covey, D. F., Whitelegge, J. P., and Evers, A. S. (2017) Photoaffinity labeling with cholesterol analogues precisely maps a cholesterol-binding site in voltage-dependent anion channel-1. J Biol Chem 292, 9294–9304

35. Budelier, M. M., Cheng, W. W. L., Chen, Z. W., Bracamontes, J. R., Sugasawa, Y., Krishnan, K., Mydock-McGrane, L., Covey, D. F., and Evers, A. S. (2018) Common binding sites for cholesterol and neurosteroids on a pentameric ligand-gated ion channel. Biochim Biophys Acta Mol Cell Biol Lipids 1864, 128–136

36. Cheng, W. W. L., Chen, Z. W., Bracamontes, J. R., Budelier, M. M., Krishnan, K., Shin, D. J., Wang, C., Jiang, X., Covey, D. F., Akk, G., and Evers, A. S. (2018) Mapping two neurosteroid-modulatory sites in the prototypic pentameric ligand-gated ion channel GLIC. J Biol Chem 293, 3013–3027

37. Chang, Y., Ghansah, E., Chen, Y., Ye, J., and Weiss, D. S. (2002) Desensitization mechanism of GABA receptors revealed by single oocyte binding and receptor function. J Neurosci 22, 7982–7990

38. Abramian, A. M., Comenencia-Ortiz, E., Modgil, A., Vien, T. N., Nakamura, Y., Moore, Y. E., Maguire, J. L., Terunuma, M., Davies, P. A., and Moss, S. J. (2014) Neurosteroids promote phosphorylation and membrane insertion of extrasynaptic GABAA receptors. Proc Natl Acad Sci U S A 111, 7132–7137

39. Comenencia-Ortiz, E., Moss, S. J., and Davies, P. A. (2014) Phosphorylation of GABAA receptors influences receptor trafficking and neurosteroid actions. Psychopharmacology (Berl) 231, 3453–3465

40. Smith, S. S., Shen, H., Gong, Q. H., and Zhou, X. (2007) Neurosteroid regulation of GABA(A) receptors: Focus on the alpha4 and delta subunits. Pharmacol Ther 116, 58–76

41. Vauquelin, G., Van Liefde, I., and Swinney, D. C. (2015) Radioligand binding to intact cells as a tool for extended drug screening in a representative physiological context. Drug Discov Today Technol 17, 28–34

42. Bylund, D. B., Deupree, J. D., and Toews, M. L. (2004) Radioligand-binding methods for membrane preparations and intact cells. Methods Mol Biol 259, 1–28

43. Bylund, D. B., and Toews, M. L. (1993) Radioligand binding methods: practical guide and tips. Am J Physiol 265, L421–429

44. Germann, A. L., Pierce, S. R., Burbridge, A. B., Steinbach, J. H., and Akk, G. (2019) Steady-State Activation and Modulation of the Concatemeric alpha 1 beta 2 gamma 2L GABA(A) Receptor. Molecular Pharmacology 96, 320–329

45. Germann, A. L., Pierce, S. R., Senneff, T. C., Burbridge, A. B., Steinbach, J. H., and Akk, G. (2019) Steady-state activation and modulation of the synaptic-type alpha1beta2gamma2L GABAA receptor by combinations of physiological and clinical ligands. Physiol Rep 7, e14230

46. Eaton, M. M., Germann, A. L., Arora, R., Cao, L. Q., Gao, X., Shin, D. J., Wu, A., Chiara, D. C., Cohen, J. B., Steinbach, J. H., Evers, A. S., and Akk, G. (2016) Multiple Non-Equivalent Interfaces Mediate Direct Activation of GABAA Receptors by Propofol. Curr Neuropharmacol 14, 772–780

47. Das, J. (2011) Aliphatic diazirines as photoaffinity probes for proteins: recent developments. Chem Rev 111, 4405–4417

48. Li, G. D., Chiara, D. C., Sawyer, G. W., Husain, S. S., Olsen, R. W., and Cohen, J. B. (2006) Identification of a GABAA receptor anesthetic binding site at subunit interfaces by photolabeling with an etomidate analog. J Neurosci 26, 11599–11605

49. Jayakar, S. S., Zhou, X., Chiara, D. C., Jarava-Barrera, C., Savechenkov, P. Y., Bruzik, K. S., Tortosa, M., Miller, K. W., and Cohen, J. B. (2019) Identifying Drugs that Bind Selectively to Intersubunit General Anesthetic Sites in the alpha1beta3gamma2 GABAAR Transmembrane Domain. Mol Pharmacol 95, 615–628

50. Stewart, D., Desai, R., Cheng, Q., Liu, A., and Forman, S. A. (2008) Tryptophan mutations at azietomidate photo-incorporation sites on alpha1 or beta2 subunits enhance GABAA receptor gating and reduce etomidate modulation. Mol Pharmacol 74, 1687–1695

51. Ziemba, A. M., Szabo, A., Pierce, D. W., Haburcak, M., Stern, A. T., Nourmahnad, A., Halpin, E. S., and Forman, S. A. (2018) Alphaxalone Binds in Inner Transmembrane beta+-alpha-Interfaces of alpha1beta3gamma2 gamma-Aminobutyric Acid Type A Receptors. Anesthesiology 128, 338–351

52. Steinbach, J. H., and Akk, G. (2001) Modulation of GABA(A) receptor channel gating by pentobarbital. J Physiol 537, 715–733

53. Bracamontes, J. R., and Steinbach, J. H. (2009) Steroid interaction with a single potentiating site is sufficient to modulate GABA-A receptor function. Mol Pharmacol 75, 973–981

54. Haage, D., and Johansson, S. (1999) Neurosteroid modulation of synaptic and GABA-evoked currents in neurons from the rat medial preoptic nucleus. J Neurophysiol 82, 143–151

55. Zhu, W. J., and Vicini, S. (1997) Neurosteroid prolongs GABAA channel deactivation by altering kinetics of desensitized states. J Neurosci 17, 4022–4031

56. Bianchi, M. T., and Macdonald, R. L. (2003) Neurosteroids shift partial agonist activation of GABA(A) receptor channels from low-to high-efficacy gating patterns. J Neurosci 23, 10934–10943

57. Jones, M. V., and Westbrook, G. L. (1995) Desensitized states prolong GABAA channel responses to brief agonist pulses. Neuron 15, 181–191

58. Lange, Y., and Steck, T. L. (2016) Active membrane cholesterol as a physiological effector. Chem Phys Lipids 199, 74–93

59. Basak, S., Schmandt, N., Gicheru, Y., and Chakrapani, S. (2017) Crystal structure and dynamics of a lipid-induced potential desensitized-state of a pentameric ligand-gated channel. Elife 6

60. Tong, A., Petroff, J. T., 2nd, Hsu, F. F., Schmidpeter, P. A., Nimigean, C. M., Sharp, L., Brannigan, G., and Cheng, W. W. (2019) Direct binding of phosphatidylglycerol at specific sites modulates desensitization of a ligand-gated ion channel. Elife 8

61. Farrant, M., and Nusser, Z. (2005) Variations on an inhibitory theme: phasic and tonic activation of GABA(A) receptors. Nat Rev Neurosci 6, 215–229

62. Feng, H. J., and Forman, S. A. (2018) Comparison of alphabetadelta and alphabetagamma GABAA receptors: Allosteric modulation and identification of subunit arrangement by site-selective general anesthetics. Pharmacol Res 133, 289–300

63. Overstreet, L. S., Jones, M. V., and Westbrook, G. L. (2000) Slow desensitization regulates the availability of synaptic GABA(A) receptors. J Neurosci 20, 7914–7921

64. Harrison, N. L., Vicini, S., and Barker, J. L. (1987) A steroid anesthetic prolongs inhibitory postsynaptic currents in cultured rat hippocampal neurons. J Neurosci 7, 604–609

65. Chakrabarti, S., Qian, M., Krishnan, K., Covey, D. F., Mennerick, S., and Akk, G. (2016) Comparison of Steroid Modulation of Spontaneous Inhibitory Postsynaptic Currents in Cultured Hippocampal Neurons and Steady-State Single-Channel Currents from Heterologously Expressed alpha1beta2gamma2L GABA(A) Receptors. Mol Pharmacol 89, 399–406

66. Jones, M. V., and Westbrook, G. L. (1996) The impact of receptor desensitization on fast synaptic transmission. Trends Neurosci 19, 96–101

67. Miller, P. S., and Aricescu, A. R. (2014) Crystal structure of a human GABA_A_ receptor. Nature 512, 270–275

68. Jansen, M., Bali, M., and Akabas, M. H. (2008) Modular design of Cys-loop ligand-gated ion channels: functional 5-HT3 and GABA rho1 receptors lacking the large cytoplasmic M3M4 loop. J Gen Physiol 131, 137–146

69. Edgar, R. C. (2004) MUSCLE: a multiple sequence alignment method with reduced time and space complexity. BMC Bioinformatics 5, 113

70. Sali, A., and Blundell, T. L. (1993) Comparative protein modelling by satisfaction of spatial restraints. J Mol Biol 234, 779–815

71. Shen, M. Y., and Sali, A. (2006) Statistical potential for assessment and prediction of protein structures. Protein Sci 15, 2507–2524

72. Lee, J., Cheng, X., Swails, J. M., Yeom, M. S., Eastman, P. K., Lemkul, J. A., Wei, S., Buckner, J., Jeong, J. C., Qi, Y., Jo, S., Pande, V. S., Case, D. A., Brooks, C. L., 3rd, MacKerell, A. D., Jr., Klauda, J. B., and Im, W. (2016) CHARMM-GUI Input Generator for NAMD, GROMACS, AMBER, OpenMM, and CHARMM/OpenMM Simulations Using the CHARMM36 Additive Force Field. J Chem Theory Comput 12, 405–413

73. McGibbon, R. T., Beauchamp, K. A., Harrigan, M. P., Klein, C., Swails, J. M., Hernandez, C. X., Schwantes, C. R., Wang, L. P., Lane, T. J., and Pande, V. S. (2015) MDTraj: A Modern Open Library for the Analysis of Molecular Dynamics Trajectories. Biophys J 109, 1528–1532

74. Trott, O., and Olson, A. J. (2010) AutoDock Vina: improving the speed and accuracy of docking with a new scoring function, efficient optimization, and multithreading. J Comput Chem 31, 455–461

75. O’Boyle, N. M., Banck, M., James, C. A., Morley, C., Vandermeersch, T., and Hutchison, G. R. (2011) Open Babel: An open chemical toolbox. J Cheminform 3, 33

76. Meslamani, J. E., Andre, F., and Petitjean, M. (2009) Assessing the Geometric Diversity of Cytochrome P450 Ligand Conformers by Hierarchical Clustering with a Stop Criterion. J Chem Inf Model 49, 330–337

